# Versatile gamma-tubulin complexes contribute to the dynamic organization of MTOCs during *Drosophila* spermatogenesis

**DOI:** 10.1101/2024.03.20.585862

**Authors:** Elham Alzyoud, Dóra Németh, Viktor Vedelek, Titanilla Szögi, Viktória Petra Tóth, Mónika Krecsmarik, Edit Ábrahám, Zoltán Lipinszki, Rita Sinka

**Affiliations:** Department of Genetics, University of Szeged, Szeged, Hungary; Faculty of Science and Informatics, Doctoral School of Biology, University of Szeged, Szeged, Hungary; Department of Pathology, University of Szeged, Szeged, Hungary; Synthetic and Systems Biology Unit, Institute of Biochemistry, HUN-REN Biological Research Centre, HUN-REN, Szeged, Hungary; National Laboratory for Biotechnology, Institute of Genetics, HUN-REN Biological Research Centre, Szeged, Hungary

**Author notes:** These authors contributed equally to this work (Elham Alzyoud, Dóra Németh).

## Abstract

The initiation of microtubule formation is facilitated by γ-tubulin and γ-Tubulin Ring Complex (γ-TuRC) in various microtubule-organizing centers (MTOCs). While the heterogeneity of tissue-specific MTOCs and γ-TuRC in *Drosophila* testis has been described, their molecular composition and physiological significance are poorly understood. We investigated the testis-specific distribution and biochemical interaction of the canonical γ-TuRC proteins Grip163 and Grip84. We found that while Grip163 is present on the centrosome and basal body, Grip84 localizes to the centrosome and Golgi in spermatocytes and colocalizes with the testis-specific γ-TuRC at the basal body, apical nuclear tip, and near the elongated mitochondria after meiosis. We also show the apical nuclear tip localization of some γ-TuRC interacting partners and prove their binding to testis-specific γ-TuRC proteins. These results highlight and prove the importance of the different γ-TuRCs in organizing the diverse MTOCs present during the extensive rearrangement of cell organelles during the spermatogenesis of *Drosophila*.

## Introduction

The microtubule (MT) network in eukaryotic cells plays a crucial role in cell differentiation and division, as well as in the movement through the cilia and flagella. The spatial re-organization of the MT cytoskeleton is a fast and dynamic process that requires distinct subcellular nucleation sites called microtubule organizing centers (MTOCs) that act as templates for the MT arrays to initiate. The main MTOC in dividing cells is the centrosome, which consists of a pair of centrioles (an older mother and a newer daughter centriole), and the pericentriolar material (PCM). Apart from the centrosome, non-centrosomal microtubular organizational centers (ncMTOCs) are present in different cell types, but it is not known how these structures regulate the organization and dynamics of microtubules. The heterogeneity of ncMTOCs is reflected in the molecular composition and the cellular localization, and their function is unique to each cell type ^1^.

MTs are polar structures containing plus and minus end. The highly dynamic plus end is generally where the majority of MT polymerization and depolymerization occurs, while the minus end is often stabilized and capped ^2^. The main factor known to specifically associate with microtubule minus ends is the γ-tubulin ring complex (γ-TuRC) ^3,4^. PCM and ncMTOCs both recruit γTuRCs to the microtubule nucleation site ^5,6^. γTuRC is a multiprotein complex consisting of γ-tubulin and its associated proteins (γ-tubulin ring proteins (Grips) in Drosophila, or γ-tubulin complex proteins (GCPs) in human) that are essential for γTuRC integrity and MT nucleation activity. The conserved and essential core of the γTuRC is the small complex (γTuSC) that consists of two γ-tubulins, one copy of Grip84 (human GCP2) and Grip91 (human GCP3) in Drosophila ^7^. The large ring complex comprises multiple γ-TuSCs and additional Grips, including Grip75 (human GCP4), Grip128 (human GCP5), Grip163 (human GCP6), and Grip71 (NEDD1) ^8,9^. An additional component of the microtubule nucleation machinery is Mzt1, which can bind to both γ-TuSC and γ-TuRC in many eukaryotes^10–12^. Although Mzt1 is essential in most organisms, it exhibits testis-specific expression and binds to Grip91 and t-Grip91 in *D. melanogaster*^13,14^. It was shown that γTuRC genes are not essential for viability, but γTuRC is able to nucleate microtubule organization more efficiently than the small complex alone ^9^.

Similarly to mammals, Drosophila spermatogenesis involves a series of stage-dependent morphological changes when stem cells give rise to mature sperm. During this process, round spermatids of 10 μm in diameter elongate to become 1850 μm in length ^15^, which requires an extensive remodelling of existing cellular organelles, as well as the addition of new structures. After meiosis, each spermatid inherits a centriole that will serve a basal body around a ring-shaped PCM-like macromolecular structure called the centriole adjunct. The growth of the axoneme starts from the basal body and continues to elongate towards the spermatid tail **(Figure 1 a)**^16^. Opposite to the basal body, on one side of the nucleus, the acroblast starts to develop, which is a Golgi-derived organelle. During the later stages of elongation, the acroblast will be converted into the acrosome. Another important event that occurs after meiosis is the fusion of the mitochondria into two giant mitochondrial derivatives, called the nebenkern. These mitochondrial derivatives elongate with the axoneme during sperm maturation and accumulate paracrystalline material in one of them, which could provide structural support for the growing tail **(Figure 1 a)**^15,17^.

**Figure 1.**
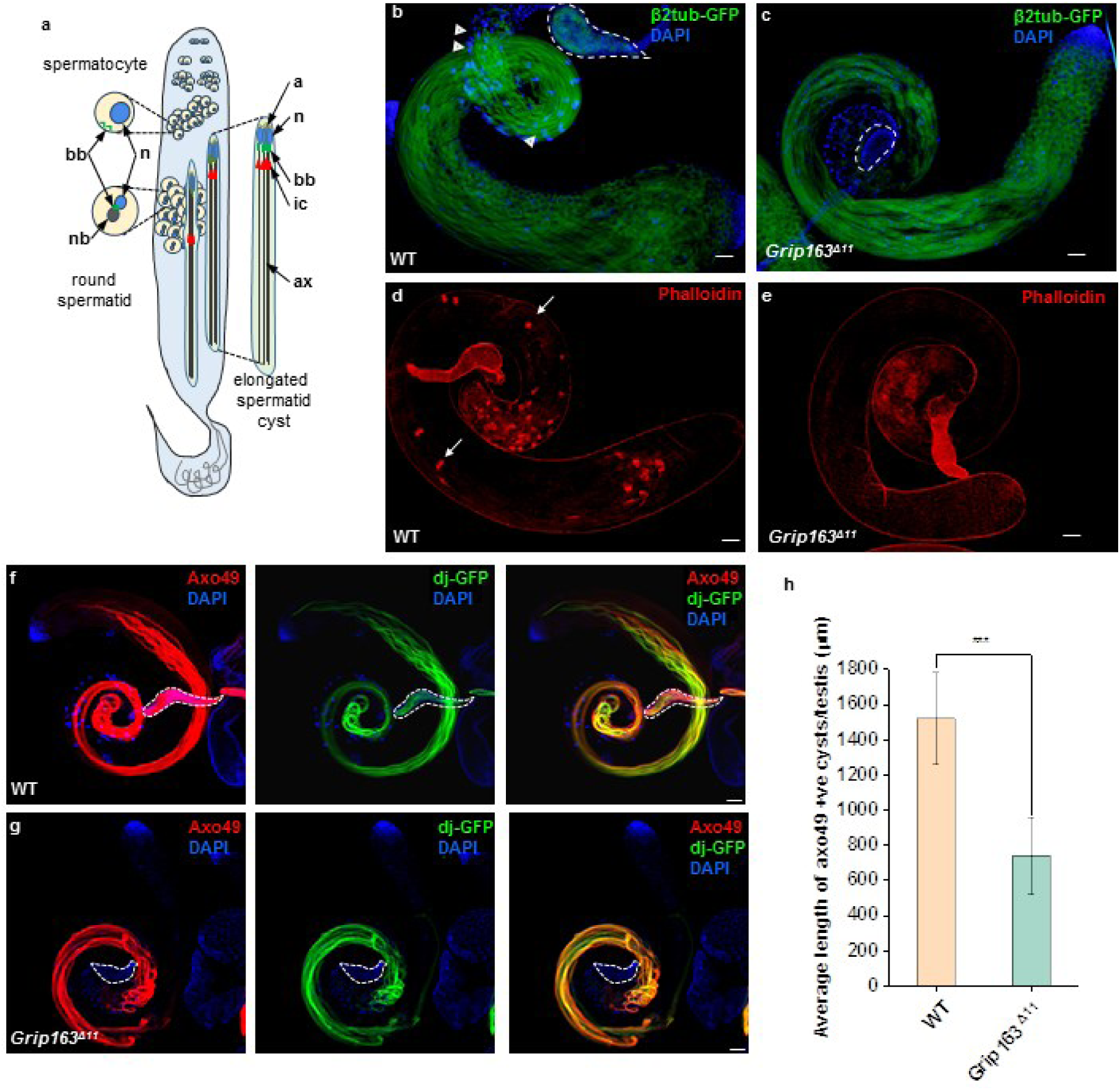
Mitochondrial and axonemal abnormalities lead to abnormal spermatid elongation in *Grip163^Δ^*^11^ *mutant*. (**a**) Schematic representation of Drosophila spermatogenesis stages in the testis, highlighting the major cellular organelles (bb: basal body, n: nucleus, nb: nebenkern, a: acrosome, ax: axoneme and cytoskeletal structures (ic: investment cone) of the secondary spermatocyte, round spermatids and the elongated spermatid cyst. **(b, c)** Microtubules are visualized with the β2tub-GFP transgene (green), while the nuclei stained with DAPI (blue) in WT **(b)** and *Grip163^Δ^*^11^ mutant spermatids **(c)** Elongated cysts are present both in WT and *Grip163^Δ^*^11^, however, the seminal vesicle (dashed lines) of *Grip163^Δ^*^11^ mutant does not contain β2tub-GFP positive mature sperms. **(d,e)** The individualization complexes (IC) were stained with Phalloidin (red), ICs are formed normally (arrows) in WT, **(c)** and missing in *Grip163^Δ^*^11^ mutant **(e)**. **(f, g)** Fully developed axonemes are visualized by Axo49 tubulin antibody (red) and elongated mitochondria are with DJ-GFP transgene (green) both in the testis and seminal vesicle (dashed line). Elongated polyglycylated axonemes and mitochondria were shorter in *Grip163^Δ^*^11^ mutant and the seminal vesicle was empty (dashed line) **(g)** compared to the WT **(f). (h)** Diagrams show the average length of Axo49 positive cysts in WT and *Grip163^Δ^*^11^ mutant. (WT n=12, *Grip163^Δ^*^11^ n=22) Statistical significance was tested by one-way ANOVA. (scale bars: 50 μm).

Growing evidence suggests that testis-specific MTOC components play important roles in basal body and ncMTOC formation and also in the positioning and shaping of the post-meiotic organelles during Drosophila spermatogenesis ^13,14,18^. For instance, testis-specific CnnT expression contributes to the conversion of mitochondria into a ncMTOC, independent of core pericentriolar Cnn ^18^. In addition to the canonical γ-TuRC genes (Grip84, Grip91, Grip75, Grip128, Grip163, and Grip71), we have recently identified three additional testis-specific paralogs of the γ-TuRC components, t-Grip84, t-Grip91, and t-Grip128 ^14,19^. We have shown, that from the round spermatid stage onward, these t-γ-TuRC proteins start to localize to the centriole adjunct, then to the nuclear tip, and finally surrounding the mitochondria of the elongating cyst. We also presented that they bind to γ-Tubulin and Mzt1 and interact with each other in the post-meiotic stages ^14^. We and others have shown that Mzt1 binds to the N-termini of Grip/GCP proteins and interacts with the N-terminal regions of Grip91, t-Grip91 and Grip128 in Drosophila ^13,14^. Our finding raised the question of whether the canonical γ-TuRC proteins are replaced with their testis-specific counterparts, or whether they are able to cooperate to meet the needs of microtubule nucleation during the late stages of spermatogenesis. We also aimed to understand whether the testis-specific γ-TuRC can interact with the canonical γ-TuRC or with any other proteins. In order to address these questions, we created *in situ* tagged *Grip163* and *Grip84* transgenic lines to elucidate the adaptable localization and identify interacting partners of the γ-TuRC during the development of male germ cells.

## Results

### Phenotypic characterization of the *Grip163 ^Δ^*^11^ mutant line

We have reported previously that t-γ-TuRC proteins are present only in the post-meiotic spermatids ^14^. However, there is a lack of data regarding the relationship between the canonical and the testis-specific γ-TuRCs, as well as the presence or localization of ubiquitously expressed γ-TuRC components in the late stages of spermatogenesis.

Therefore, we tested the presence and possible function of the γ-TuRC component Grip163, which has no testis-specific paralogs, in sperm development ^20,8,14^. To achieve this, we induced mutation in the *Grip163* gene, and performed an *in situ* tagging of the *Grip163* gene, using the CRISPR-Cas9 gene editing technology. Using a specific guided RNA designed for *Grip163*, we deleted 11 base pairs from the coding region of the gene (hereafter, *Grip163^Δ^*^11^). This deletion caused a frameshift leading to a premature stop codon in the coding region of *Grip163* (**S.Figure 1a**). Interestingly, we found that in contrast to the G*rip84* and *Grip91* homozygotes, which are lethal ^21,22^, the homozygous *Grip163^Δ^*^11^ mutant animals are viable, but male and female sterile (**S.Figure 1c**). It is known that the γ-TuSC is essential for centrosome formation, mutants of the canonically expressed γ-TuRC components, *Grip128* and *Grip75,* are viable and similar to *Grip163^Δ^*^11^ male and female sterile ^23,9,21,24,25^. The previously characterised *Dgrip163^GE^*^27087^ mutant allele is male fertile and female sterile^9^, therefore we decided to test the spermatogenesis phenotype of *Grip163^Δ^*^11^. *Grip163^Δ^*^11^ is male and female sterile in hemizygous combination with the overlapping *Df(3L)BSC574* deficiency, suggests that *Grip163^Δ^*^11^ is a strong hypomorph or a null allele (**S.Figure 1c)**. The seminal vesicles of the *Grip163^Δ^*^11^ males were empty, without mature sperms (**Figure 1 b-c**). Based on this phenotype we investigated both the early and late stages of spermatogenesis in *Grip163^Δ^*^11^ flies. We found an abnormal nebenkern-nucleus ratio in the round spermatids, which is a hallmark of abnormal mitotic division (**Figure 2 a-c)**. Nevertheless, many of the cysts are able to elongate in the *Grip163^Δ^*^11^ mutant, visualized by the β2tub-GFP transgene (**Figure 1 b, c)**. To further investigate the elongation of the cysts, we tested whether the individualization proceeds normally. After the elongation of the cyst, an actin rich cytoskeletal structure, the individualization complex (IC) is formed at the caudal end of the nuclei surrounding the basal-body and axoneme. During individualization the IC moves towards the tip of the tail, expelling and degrading all the cytoplasmic content from the spermatid and tightly cover the axoneme-mitochondria complex with an individual plasma membrane **(Figure 1 a)**^26^. To visualize the elongated post-meiotic cysts, we stained the actin cones of the IC in the *Grip163^Δ^*^11^ testis and found the lack of them in *Grip163^Δ^*^11^, providing evidence for the failure of proper cyst elongation, which is the prerequisite of individualisation **(Figure 1 d, e).** For further evidence, we used anti-pan polyglycylated tubulin (Axo49) antibody to visualize mature axonemes of the spermatids, at the onset of individualization^27^. We observed significantly shorter axonemes in the *Grip163^Δ^*^11^ mutant compared to the wild-type (WT), indicating abnormal spermatid elongation **(Figure 1 f-h).**

**Figure 2:**
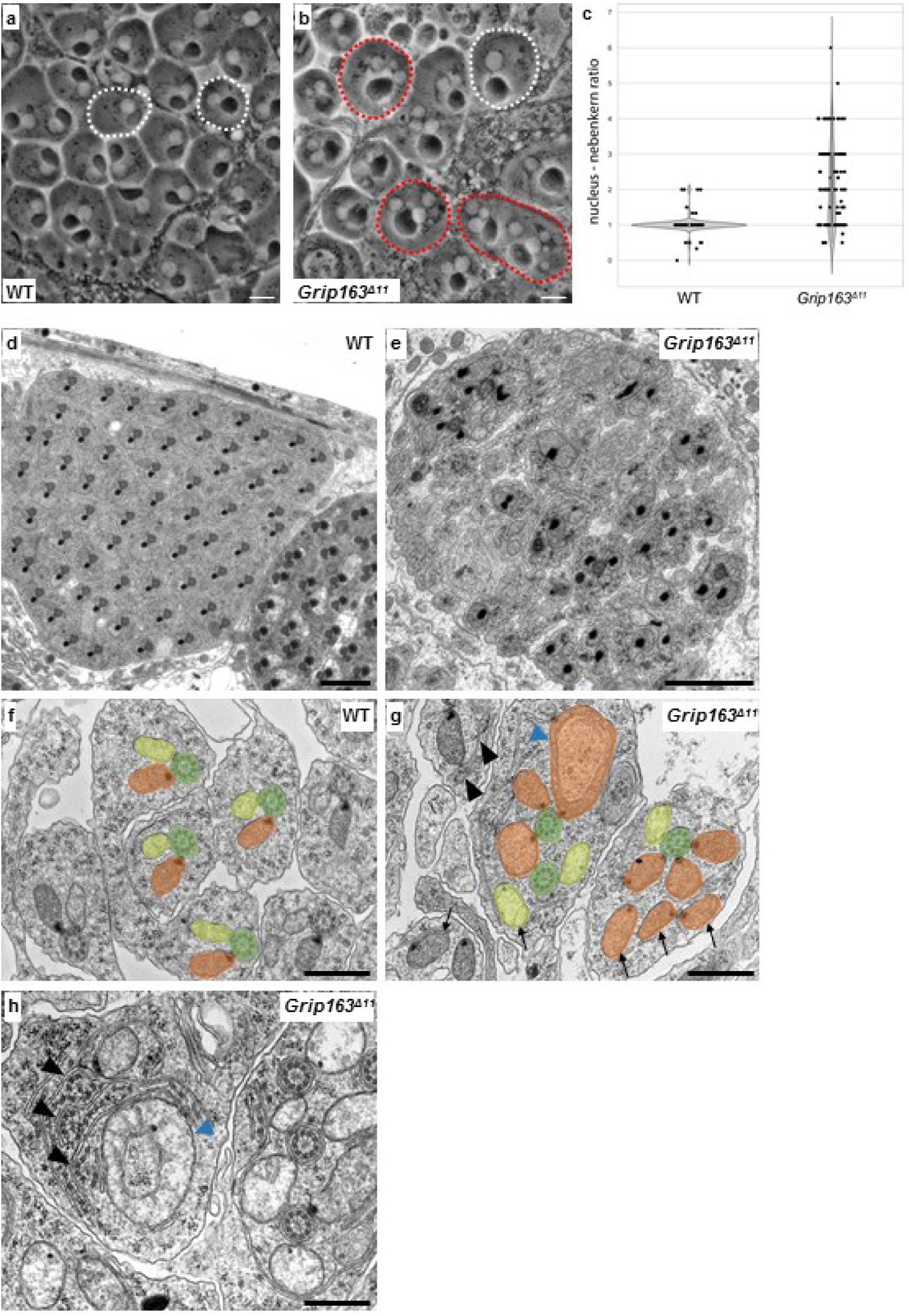
Abnormal mitochondrial and axonemal structures in the post-meiotic cysts of *Grip163^Δ^*^11^ mutant. (**a, b)** Phase contrast images show a normal 1:1 (white outline), nucleus: nebenkern ratio in WT **(a)** and an abnormal ratio (red outline) in *Grip163^Δ^*^11^ mutant **(b)**. **(c)** Violin plot showing the nucleus: nebenkern ratio in WT (n=186) and *Grip163^Δ^*^11^ mutant round spermatids (n=112). **(d, h)** Transmission electron micrograms of testis cross-sections, representing elongating cysts of WT (**d,f**) and *Grip163^Δ^*^11^ mutant (**e,g,h**). Disorganized cysts with axonemal and mitochondrial abnormalities are characteristic in *Grip163^Δ^*^11^ mutant**. (f)** WT elongating cyst shows normal axoneme structure with 9+2 tubulin dimers (highlighted in green) and two mitochondrial derivatives (yellow and orange), with paracrystalline material formation in one of them (orange). **(g, h)** In *Grip163^Δ^*^11^ mutant the mitochondria and the axoneme detached from each other, and multiple mitochondrial derivatives (highlighted in orange and yellow) attached to one axoneme (green). **(h)** Mitochondria fused, enlarged (blue arrowheads) and detached (arrow) from the axoneme and disintegrated axonemes with microtubule bundles and the axial membrane opening is characteristic of *Grip163^Δ^*^11^ mutant cyst (black arrowheads). (scale bars: A-E 2 μm, F-H 500 nm)

Axoneme elongation depends on the mitochondria, which provide a structural platform for MT reorganization to support the robust elongation taking place at the tail tip of the sperm^28^. Based on this we tested the presence and appearance of late elongating mitochondria using dj-GFP as a marker. In accordance with the defect observed in axoneme elongation, it was detected that the mitochondria in the *Grip163^Δ^*^11^ mutants also exhibited incomplete elongation (**Figure 1 f-g**).

Based on this general spermatid elongation defect of *Grip163^Δ^*^11^ flies, the detailed morphology of post-meiotic cysts was further investigated using transmission electron microscopy (TEM). In wild-type testis, the 64 spermatids develop simultaneously and on the TEM image of elongated cysts we observed the two mitochondrial derivatives attached to the axoneme, which has the classical 9+2 arrangement of microtubules in each spermatid (**Figure 2 d-f**). In *Grip163^Δ^*^11^ mutants, cysts became disorganized, with fewer spermatids per cyst and with disturbed mitochondrial:axonemal ratio (**Figure 2. e**). Frequently, multiple mitochondrial derivatives were attached to a single axoneme **(Figure 2 g)**, and we also observed multiple damaged axonemes **(Figure 2 h)**, supporting that Grip163 is necessary for normal axoneme formation and/or stability. Since the γ-TuRC was proposed to be essential for centrosome formation in somatic cells ^11^, we examined the centrosome/basal body distribution at different stages of spermatogenesis in the testes of *Grip163^Δ^*^11^ males. We tested the presence and the distribution of the centrosomal/basal body localized γ-Tubulin and the PACT domain of DPlp fused with GFP (GFP-PACT)^29^. In the wild-type animals both proteins localize to the centrosome/basal body during different stages of spermatogenesis **(Figure 3 a-b, S**.**Figure 2b)**. In wild-type males, GFP-PACT is located close to the nucleus and the nebenkern, as expected in round spermatids. In contrast, multiple and scattered GFP-PACT signals were observed in the *Grip163^Δ^*^11^ mutants’ testis (**Figure 3 c-d, S**.**Figure 2b-c)**. We calculated both the nucleus-nebenkern and the basal body-nebenkern ratio in the round spermatids **(S.Figure 2a)**. Theoretically, these ratios should be close to 1 in WT, as each spermatid is supposed to have one of each organelles per cell. However, we found that the *Grip163^Δ^*^11^ mutant were significantly different from the expected 1. In the *Grip163^Δ^*^11^ mutant, the average ratio is higher than one (∼2.06, ∼2.01) indicating that there are less nebenkerns but approximately equal amount of basal bodies and nuclei **(S.Figure 2a)**. These abnormalities could be the consequence of cytokinetic defects of the meiotic cysts of *Grip163^Δ^*^11^ mutant.

**Figure 3:**
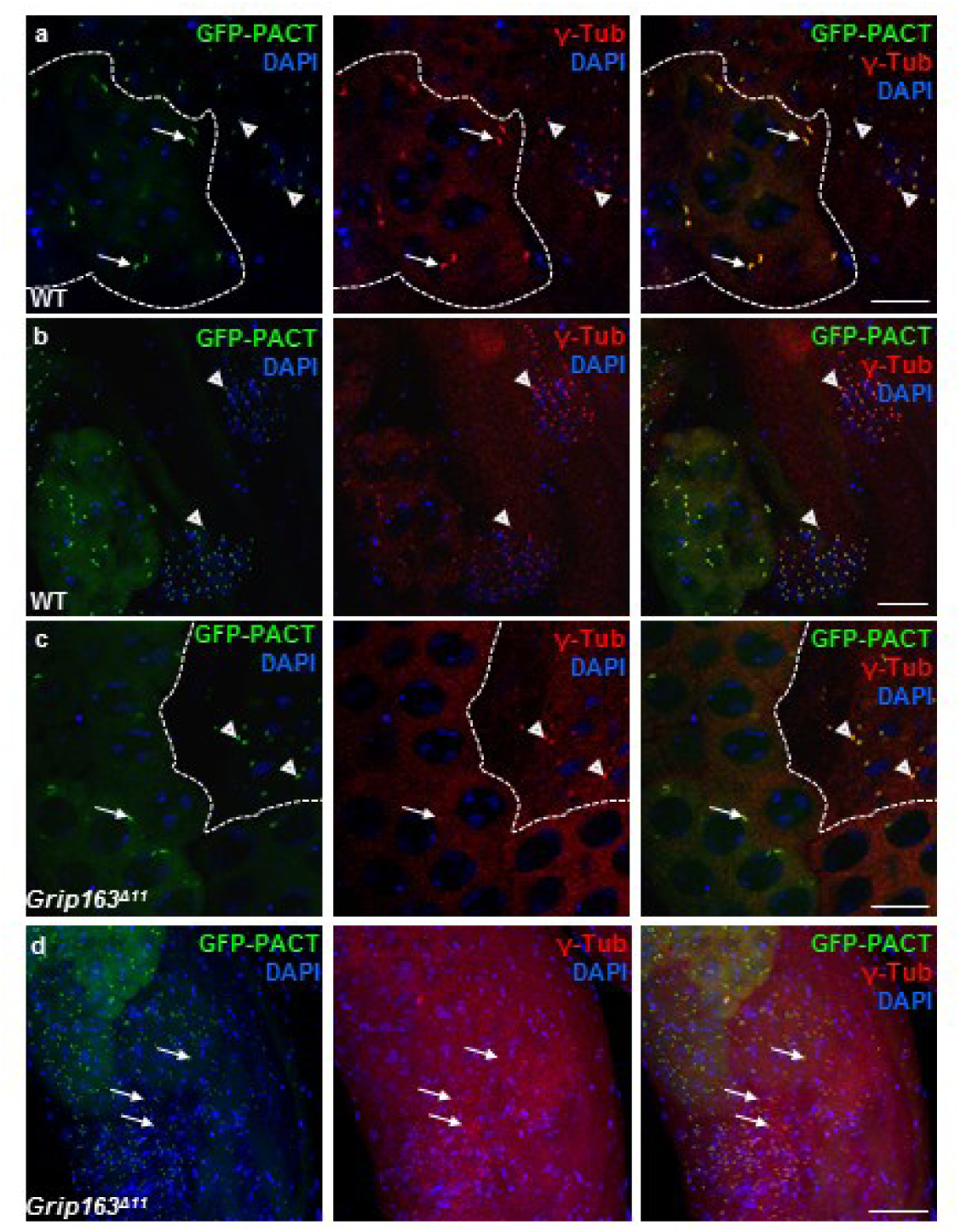
Mislocalization of centrosomal markers in *Grip163^Δ11^*mutant. **(a-b)** GFP-PACT (green) and γ-Tubulin (red) are colocalizing on the centrosome (arrows) and basal body (arrowheads) in WT spermatocytes and spermatids **(a, b)**. γ-tubulin, but not GFP-PACT centrosomal localization is affected in *Grip163^Δ^*^11^ mutant spermatocytes. In the round spermatids, GFP-PACT and γ-Tubulin are colocalised **(c)**, but the post-meiotic elongating cysts are disorganized with scattered basal bodies. **(d)**. (scale bars: 20 μm)

We also measured the fluorescence intensity of γ-Tubulin and GFP-PACT in premeiotic and post-meiotic cysts. Both in WT and the *Grip163^Δ^*^11^ mutant flies we found strong GFP-PACT positive centrosomal and basal body signals **(Figure 3 c-d)**, however, the γ-Tubulin signal was weak in spermatocytes compared to the WT spermatocytes **(S.Figure 2d-e)**.

Surprisingly, after meiosis the γ-Tubulin signal was stronger and accumulated on the basal body both in WT and *Grip163^Δ^*^11^ mutant spermatids, but those were scattered along the cysts (**Figure 3 c, d**, **S.Figure 2e)**. It was previously proposed that γ-Tubulin could be recruited to the centrosomes in somatic cells by γ-TuSCs, independently of the γ-TuRCs ^9^. Mzt1 is a well characterised binding partner of the γ-TuSC and we showed previously, that the post-meiotically localizing three t-Grip proteins (t-Grip84, t-Grip91, t-Grip128) bind to γ-Tubulin and t-Grip91 and also to Mzt1 ^14,30^. The γ-TuRC interacting Mzt1 is localised to the centrosome, basal body and the surface of the mitochondria from the meiotic stages onward and also to the apical tip of the elongating nuclei **(Figure 9)** ^14^. We found Mzt1 localization on the surface of the mitochondria in *Grip163^Δ^*^11^ mutant, suggesting that Grip163 and probably γ-TuRC are not essential for Mzt1 mitochondrial anchoring. Similar to γ-Tubulin, Mzt1 is not present on the meiotic basal body, however, in the post-meiotic stages, it localizes to the dispersed basal body in the *Grip163^Δ^*^11^ mutant spermatids **(S.Figure 3).** All these data suggest that Grip163 could be essential for the formation of stable γ-TuRC before and at the onset of meiosis, however, the newly organized testis-specific γ-TuRC could contribute to the γ-Tubulin and Mzt1 binding after meiosis **(Figure 9).**

### Localization of γ-TuRC proteins during spermatogenesis

To test the subcellular localization of Grip163 at different stages of spermatogenesis, we generated endogenously GFP-tagged Grip163 transgenic lines. We and others have shown that the N-or C-termini of the γ-TuRC proteins are responsible for the binding of their interacting partners ^13,14^, therefore we tagged the Grip163 close the N-terminus (aa150) (**S.Figure 1)**, without interfering with the functional GCP domains of the protein (**S.Figure 1 a**). The fertility of the GFP-Grip163 males is comparable to the wild-type animals, indicating that the inserted fluorescent tag does not alter the function of the protein (**S.Figure 1 c)**. GFP-Grip163 signal is present and colocalised with γ-Tubulin on the centrosomes and basal body on the developing spermatids **(Figure 4 a-b)**.

**Figure 4:**
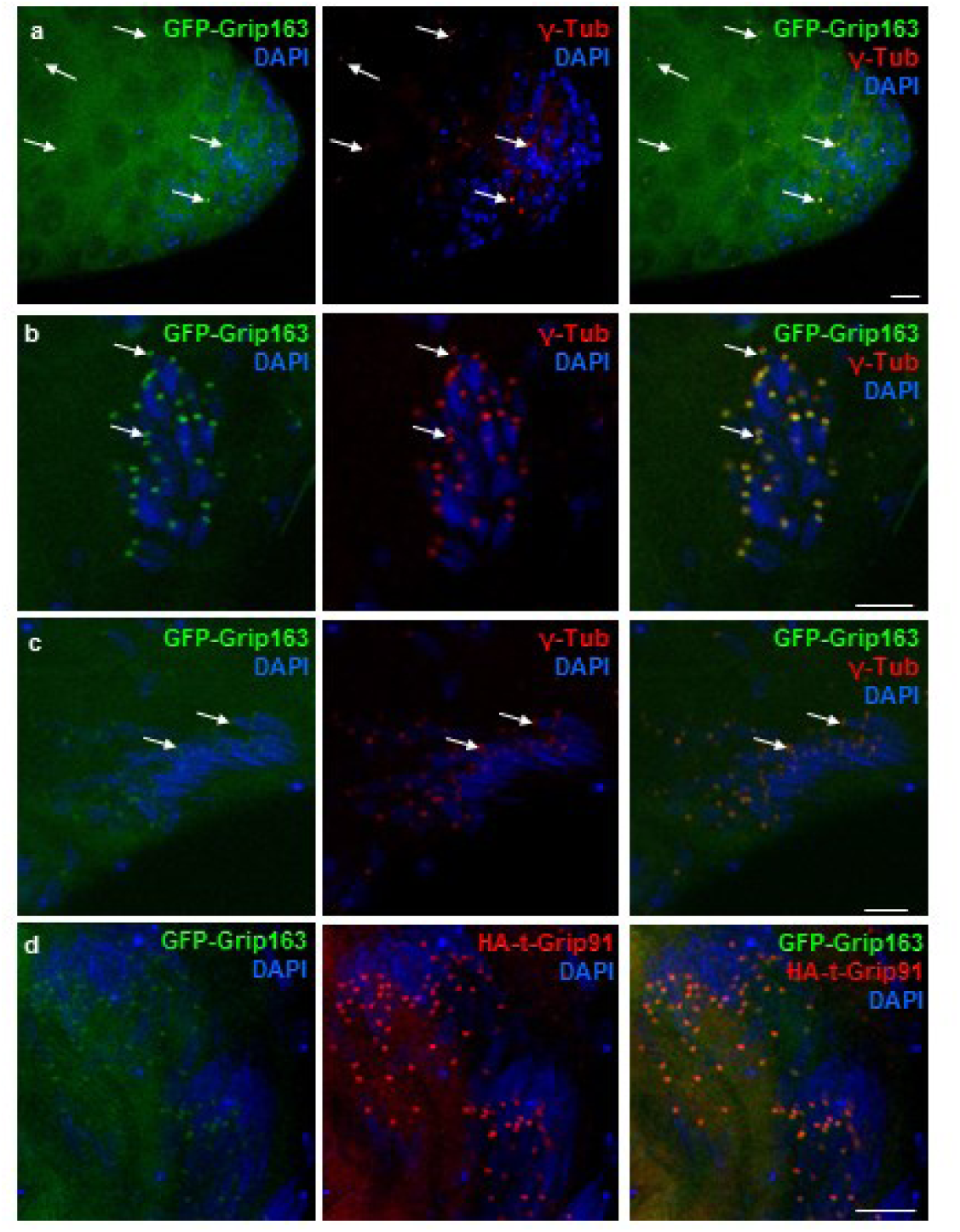
Localization of GFP-Grip163 during spermatogenesis. **(a-c)** The endogenously tagged GFP-Grip163 colocalizes with γ-Tubulin at the **(a)** centrosome (arrows) and the **(b-c)** basal body (arrows) throughout spermatogenesis. **(d)** GFP-Grip163 has an overlapping localization with the t-γ-TuRC represented by HA-t-Grip91 on the basal body/centriole adjunct in the elongating spermatids). (scale bars: 10 μm)

We also showed the colocalization of GFP-Grip163 with γ-Tubulin and also with HA-t-Grip91 on the post-meiotic basal body/centriole adjunct **(Figure 4 c-d)**. These results strongly suggest that the testis-specific γ-TuRC does not replace the canonical γ-TuRC, but both are present on the basal body/centriole adjunct of the developing spermatids **(Figure 9)**.

The localization of GFP-Grip163 to the post-meiotic centriole adjunct has prompted an inquiry into the potential presence of other canonical γ-TuSC components in the subsequent developmental stages. To investigate this, we applied CRISPR technology to *in situ* tag Grip84 with GFP. We designed the incorporation in a way that all 5 potential isoforms of Grip84 will have the N-terminally inserted GFP tag, outside of the GCP domain (**S.Figure 1 b**). It was previously reported that the classical mutant of *Grip84* is lethal, therefore the normal viability and fertility of the *in situ* GFP-tagged Grip84 serves as evidence that the GFP tag does not alter the normal function of Grip84 (**S.Figure 1 c**) ^31^. We found that GFP-Grip84 is present on the centrosomes of spermatocytes, similarly to the GFP-Grip163 protein (**Figure 5 a-e**). This localization of GFP-Grip84 was also verified by its colocalization with the centrosomal component Asterless (Asl) (**Figure 6 a, S.Figure 4 a, b, Figure 9**) ^32^.

**Figure 5:**
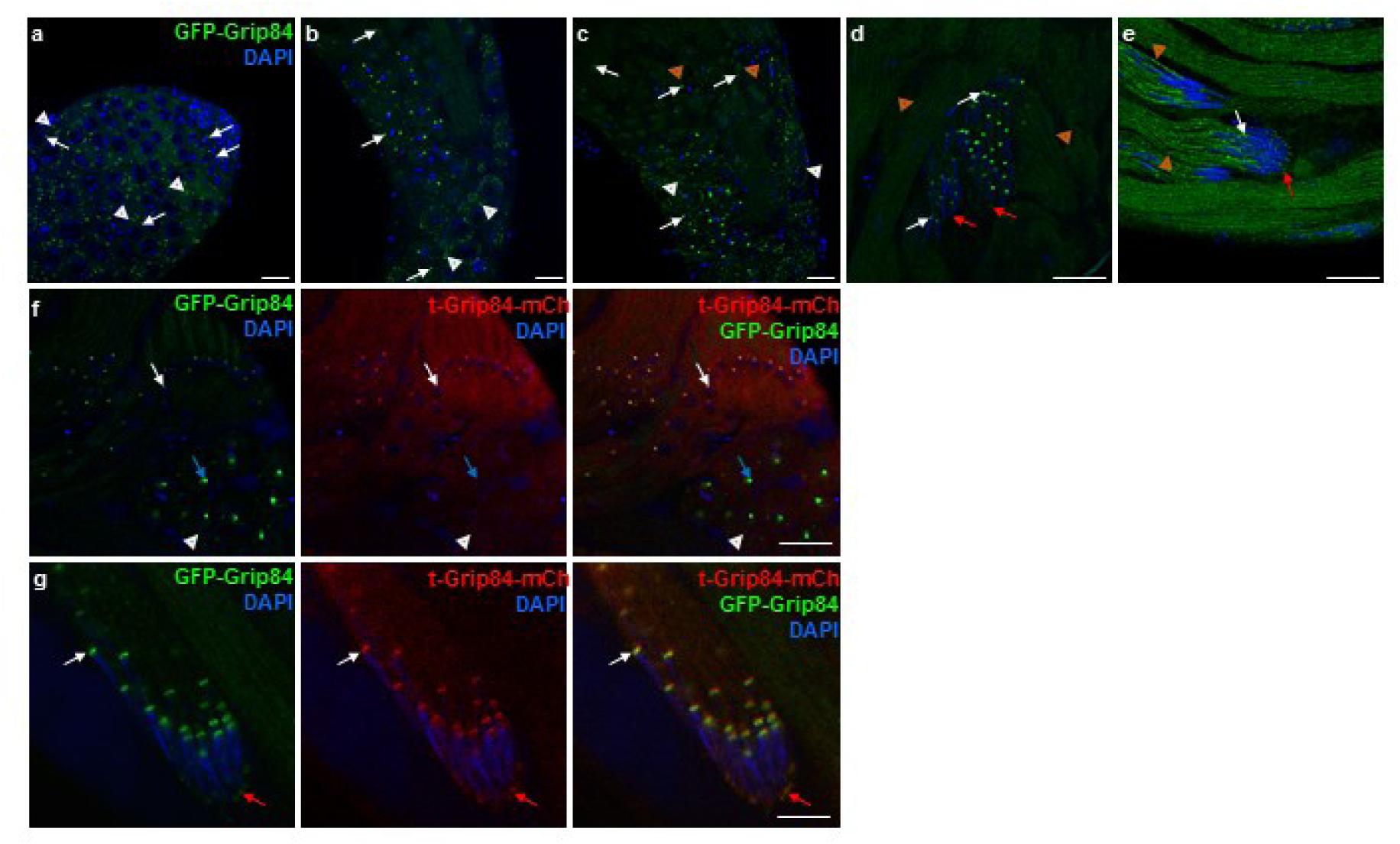
Localization of GFP-Grip84 during spermatogenesis. **(a-e)** *In vivo* tagged GFP-Grip84 localizes both to the centrosome (**a**) and the basal body (**b**, **c**) (white arrows) and in the post-meiotic elongating cyst GFP-Grip84 localizes also to the apical tip of the elongating nuclei (**d, e**) (red arrows). Additionally, it is localized surrounding the elongating mitochondria (**c-e**) (orange arrowheads). Golgi-like localization of GFP-Grip84 was observed in spermatocytes (**c**) (white arrowheads). **(f, g)** GFP-Grip84 localizes to the centrosome before meiosis (blue arrows) and shows overlapping localization with t-Grip84-mCh at the basal body (white arrows) and the apical tip of the elongating nuclei (red arrows) in the post-meiotic spermatids (Scale bars: 20 μm).

**Figure 6:**
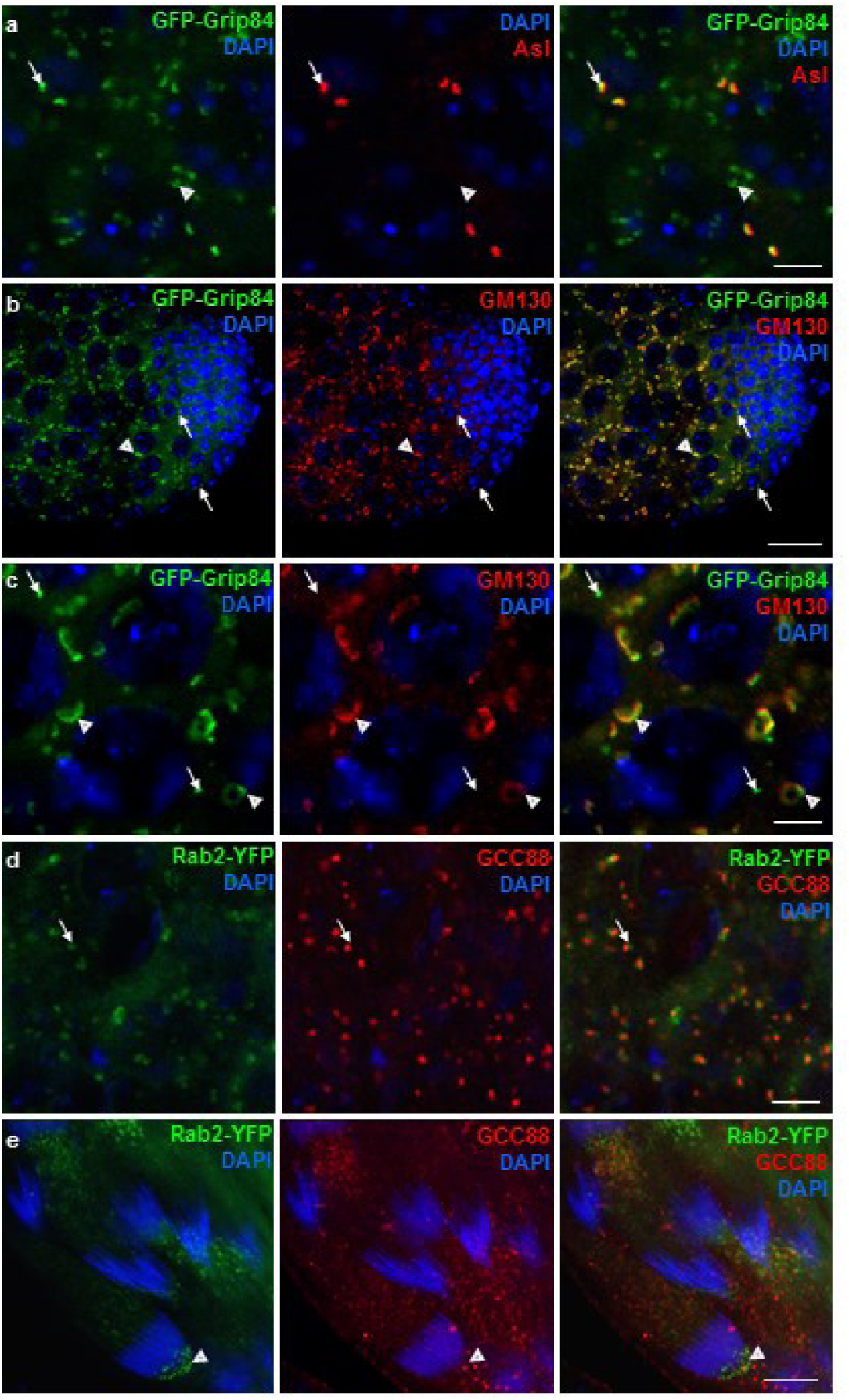
γ-TuSC protein GFP-Grip84 is localizing to the Golgi in spermatocytes. **(a)** GFP-Grip84 colocalizes with centrosomal protein Asl (anti-Asl, white arrow) and also to the Golgi apparatus (white arrowhead). **(b, c)** GFP-Grip84 colocalizes with antibody-stained *cis*-Golgi marker GM130 in the spermatocytes (arrowheads). **(d, e)** Rab2-YFP localizes to the Golgi (arrow) in spermatocytes visualised by the *trans-*Golgi marker GCC88 and also to the apical end of the elongating nuclei (arrowhead). (Scale bars: A, C, D 10 μm, B, E20 μm).

Following meiosis, GFP-Grip84 is present on the centriole adjunct and then appears at the apical nuclear tip and exhibits a strong overlap with the elongated mitochondria, akin to the GFP-Grip163. We wanted to know whether the post-meiotic localization of GFP-Grip84 overlaps with its testis-specific paralogue, t-Grip84. To do this we co-expressed GFP-Grip84 and t-Grip84-mCherry together and found complete colocalization after the meiotic stages (**Figure 5 f, g**) ^14^. These results provide additional evidence that both the ubiquitously expressed γ-TuRC and the testis-specific γ-TuRC may collectively contribute to the formation of the diverse microtubule organizing centers during the post-meiotic stages of spermatids. Apart from the centrosomal localization of GFP-Grip84, we also noticed strong bright punctae in the cytoplasm of spermatocytes, whose distribution is reminiscent of t-ER-Golgi units. It is known that centrosomal proteins are associated with the Golgi apparatus in mammalian cells ^33^. Golgi organization is special in Drosophila, as the scattered Golgi stacks are found in close proximity to the ER exit sites (t-ER-Golgi) ^34^. To test the t-ER-Golgi localization of GFP-Grip84, we used GM130 as a *cis-*Golgi, as well as GCC88, as a *trans-*Golgi-specific antibody for co-immunostaining (**Figure 6 b-e, S**.**Figure 4 c-d**, **Figure 9**). We observed strong co-localization of GFP-Grip84 with the *cis-*Golgi, suggesting that GFP-Grip84 is connected to the t-ER-Golgi units of spermatocytes.

In a comprehensive analysis of the effector proteins of Rab GTPases, Grip84, Grip91 and γ-Tubulin were identified as interacting partners of the Golgi-localizing Rab2 and Rab4 ^35^. Rab2A and Rab2B proteins were shown to be essential for acrosome formation and male fertility in mice ^36^, therefore it seemed reasonable to check whether the Golgi-localizing Rab2 might also be present at the apical end where the acrosome is located during nuclear elongation in Drosophila spermatids. We tested the germline-specific Bam-Gal4 driven pUASP-Rab2-YFP localization in spermatocytes and spermatids. We detected Rab2-YFP on the t-ER-Golgi units adjacent to the *trans-*Golgi-specific antibody GCC88 in spermatocytes and on the acrosome containing apical nuclear tip of the elongated spermatids (**Figure 6 d, e**). Localization of GFP-Grip84 to the Golgi apparatus led us to hypothesize that Drosophila γ-TuRC, similarly to the mammalian γ-TuRC might be necessary to the Golgi organization and also could have a possible function of Rab2 in acrosome formation **(Figure 9)**. Additional experiments can address these highly intriguing prospects.

### Post-meiotic ncMTOCs contain centrosomal components

The versatile localization pattern of GFP-Grip84 prompted us to investigate whether any of the already known binding partners of the γ-TuRC could be present on the newly identified localization foci. It was shown previously that γ-Tubulin is localized in the apical region of the spermatid nuclei, specifically at the distal end of the MTs present within the dense complex ^14,37^. We have previously demonstrated the presence of t-γ-TuRC and Mtz1 at the apical tip of the nucleus, suggesting the presence of a ncMTOC organized opposite to the basal body during nuclear elongation ^14^. Therefore, we carefully examined the precise localization of the centriole component, Sas4, a known interacting partner of γ-TuRC proteins by immunostaining using anti-Sas4 antibody ^38,39^. We detected Sas4 on the centrosome of mitotic spermatocytes and the basal body of the spermatids after meiosis (**Figure 7 a-c**). Intriguingly, we detected endogenous Sas4 on the apical tip of the elongating spermatid nuclei as well (**Figure 7 c**). Apart from the γ-TuRC proteins, Sas4 has multiple centriolar binding partners, including Sas6, Asl, Ana1, and Dplp ^32,39–42^. Therefore, we wondered whether they are also present on the site, which has been suggested as a ncMTOC at the apical tip of the spermatid nuclei. To achieve this we performed immunostainings using gene-specific antibodies ^32,38,43^.

**Figure 7:**
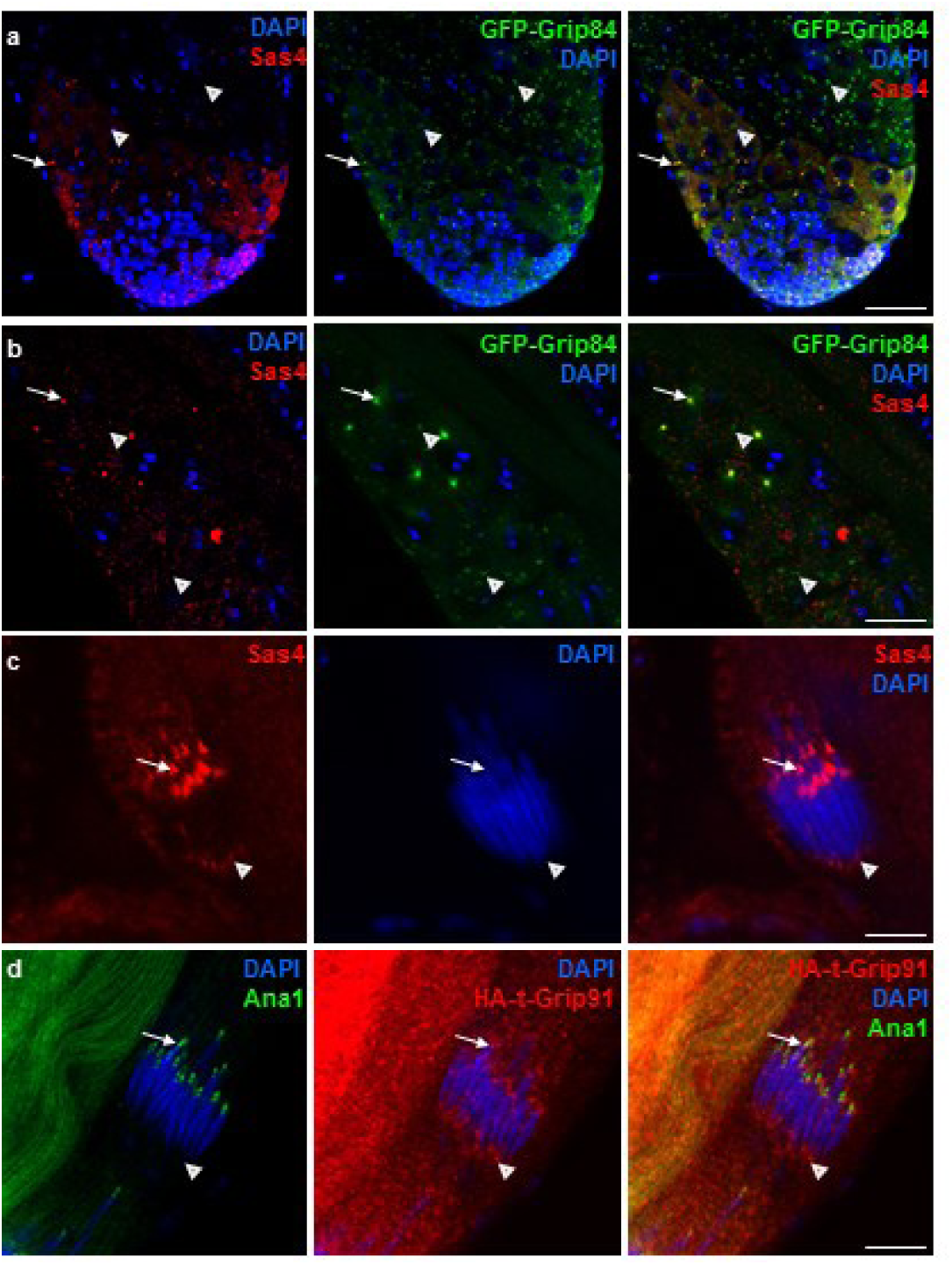
Sas4 is a component of the site, which has been suggested as a ncMTOC at the apical tip of the elongating nuclei. **(a, b)** Antibody-stained Sas4 (red) colocalizes with the endogenous tag GFP-Grip84 at the centrosome (arrows) but not on the Golgi (arrowheads) in the spermatocytes. **(c)** Sas4 (red) is localizing to the basal body of the elongating spermatids (arrows) and additionally to the apical tip of the nuclei (arrowheads). **(d)** Antibody-stained Ana1 localizes only to the basal body (arrows) of the elongating spermatids, while the t-γ-TuRC component HA-t-Grip91 localizes to the basal body(arrow) and the apical tip (arrowheads) of the elongating spermatids. (Scale bars: 20 μm).

We detected Sas6 and Asl at the centriole adjunct and also at the apical tip of the nuclei (**S.Figure 5 a, b**), however, Ana1 and GFP-PACT colocalized with t-Grip91 exclusively at the centriole adjunct region of the elongating leaf and canoe stage nuclei, in line with the reported localization pattern of these proteins (**Figure 7 d, S.Figure 5 c**) ^37^. These results indicate that the site which has been suggested as a ncMTOC at the apical nuclear tip contains several well-defined binding partners of the γ-TuRC **(Figure 9)**.

### Composition of t-γ-TuRC

We next examined whether the partially overlapping localization pattern might be a consequence of the physical interaction between the canonical γ-TuRC and the t-γ-TuRC and/or the t-γ-TuRC proteins and the γ-TuRC interacting protein Sas4. To address this, we performed a yeast two-hybrid (Y2H) analysis, where we tested t-γ-TuRC proteins with the canonical γ-TuRC protein Grip91, Grip84, Grip163, Grip75 and Sas4. Due to the large size of these proteins, we split all three t-γ-TuRC proteins as well as Grip91, Grip163 and Grip128 into an N-terminal (NT) and a C-terminal (CT) part. Grip84, Grip75 and Sas4 were used as full-length **(S.Figure 6a)**. A specific interaction was found between t-Grip84-CT and Grip91-NT, which suggests the possibility of the presence of a testis-specific and non-testis-specific mixed γ-TuRC (**Figure 8 a, S.Figure 6-7**). We also detected interaction between t-Grip84-CT and Grip84, between t-Grip91-NT and Grip84, Grip91-CT, Grip75, Grip163-NT, Grip163-CT and finally between t-Grip91-CT and Grip84, Grip91-CT, Grip75, Grip163-NT, Grip163-CT. Sas4 was able to bind to and t-Grip91-NT, t-Grip91-C, t-Grip128-N and t-Grip128-C however, we could not find an interaction between Sas4 and t-Grip84-N, t-Grip84-C (**Figure 8 a, S.Figure 6-7**).

**Figure 8:**
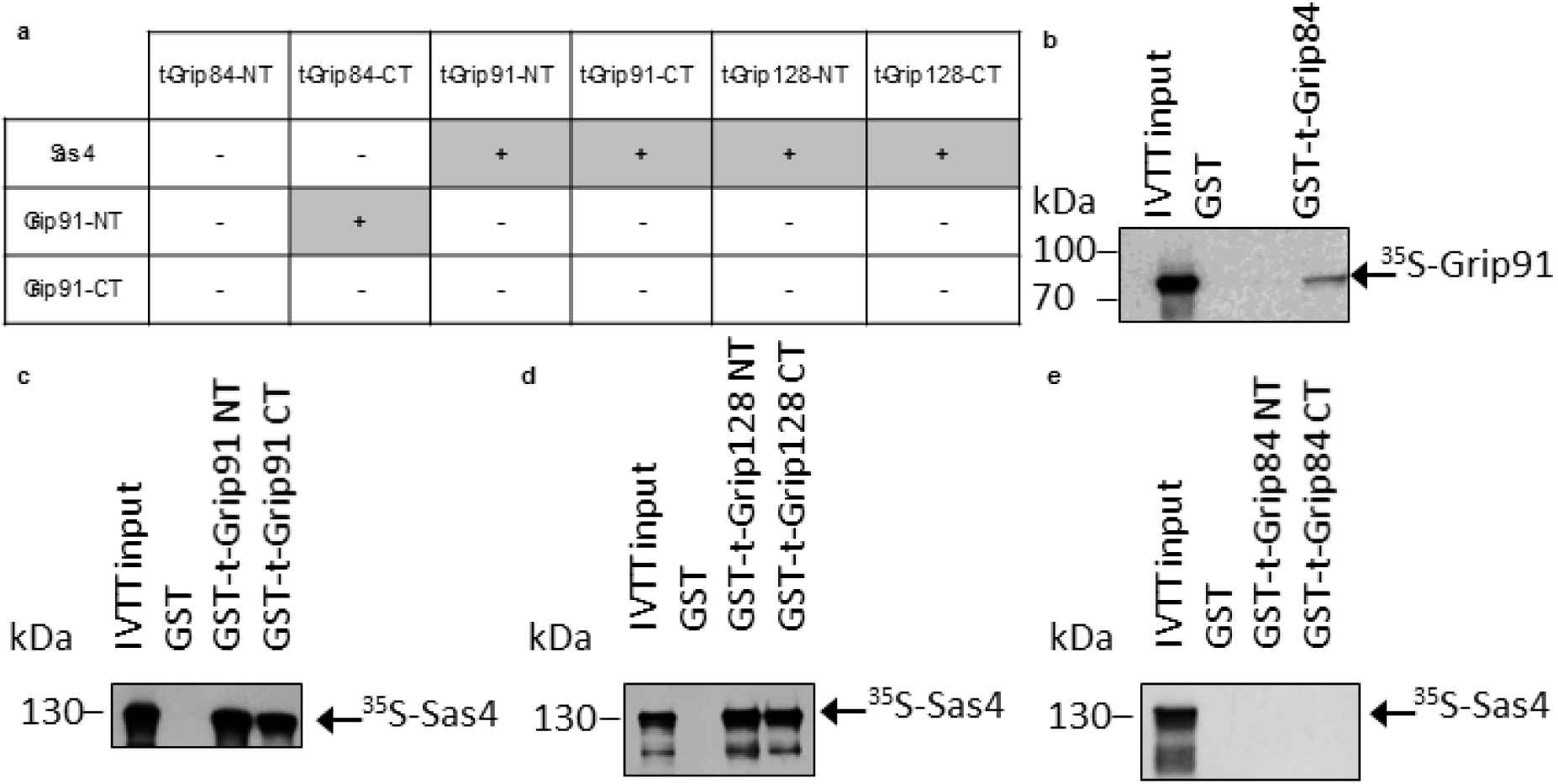
Interactions of t-γ-TuRC proteins with Sas4 and the canonical Grip91. **(a)** Summary of Y2H analysis of the N- or C-terminal parts of t-γ-TuRC proteins (as indicated in the table) with Sas4, and the N- or C-terminal parts of Grip91, respectively. **(b-e)** Autoradiographic images show the presence or absence of interaction between purified and immobilised GST-t-γ-TuRC protein fragments and ^35^S-labelled Sas4 or Grip91, synthesized by *in vitro* transcription-translation (IVTT) reaction.

To support Y2H results and prove the direct interaction between proteins, we performed an *in vitro* binding experiment and tested several combinations of proteins that showed interaction in the Y2H assay. We used immobilised GST-tagged recombinant proteins (N- or C-terminal parts of the t-γ-TuRC proteins, as indicated) as bait, which were mixed with ^35^S-methionine-labelled Sas4 or Grip91, respectively, produced by a coupled *in vitro* transcription-translation (IVTT) reaction followed by autoradiography (**Figure 8 b-e, S.Figure8**). The *in vitro* binding experiments confirmed the results of the Y2H assay (**Figure 8 a-e**). Both the localisation and the interaction studies suggest the presence of different γ-TuRC/t-γ-TuRC complexes coexisting. To further investigate these complexes, we conducted biochemical isolation and analysis of the subunit composition and migration pattern of various γ-tubulin-containing complexes in testis samples. This involved fractionating protein complexes from testes extracts using Superose 6 size-exclusion chromatography (SEC) ^44^. As a control, we fractionated canonical γ-Tubulin complex components from embryo extract (0-5 hours embryos contain somatic centrosomal MTOCs with canonical γ-TuRC, but without t-γ-TuRC proteins) of the γ-TuSC component GFP-Grip84 and γ-TuRC component GFP-Grip163, and probed them with anti-GFP, anti-γ-Tubulin and anti-p54 antibodies **(S.Figure 9)**. To estimate the relative molecular weights of the analysed protein complexes, we compare them to the SEC profile of p54/Rpn10, the ubiquitin receptor subunit of the 2,5 MDa 26S Proteasome^45^. The same extraction and fractionation method was used to analyse testes extracts by SEC, and to investigate the presence of different t-γ-TuRC in each fraction. We analysed GFP-Grip163 and GFP-Grip84, γ-Tubulin and members of the testis-specific complexes, HA-t-Grip91 and t-Grip84-mCh. Their SEC profiles suggest that they co-migrate in the high molecular weight fractions, and partially co-migrate (their peaks partially overlap) in the low molecular weight fractions **(S.Figure 9)**.

## Discussion

Drosophila spermiogenesis and spermatogenesis represent an ideal model for studying mitotic and meiotic cell division, centrosome and basal body formation and dynamic organelle reorganization after meiosis. It was proven to be a good model for understanding centrosomal and non-centrosomal MT nucleation due to the intensive morphological changes and organelle movement in the post-meiotic spermatids ^13,14,18^. MT nucleation was reported, among others, at the surface of the Golgi ^46^, mitochondria ^18^, nuclear envelope ^47^ and plasma membrane ^48^ in mammalian cells. We previously identified and described the function and versatile localization of three testis-specific γ-TuRC proteins in the post-meiotic spermatids ^14^(**Figure 9**). The described centriole adjunct, nuclear tip and mitochondrial surface localization of the t-γ-TuRC proteins and their interaction partner Mzt1 raised the question of whether the ubiquitously expressed γ-TuRC is replaced after meiosis for the testis-specific one or both are present due to the multiple putative ncMTOC formation during spermatid elongation and acrosome formation. γ-Tubulin was already reported to localize to both tips of the elongating nuclei and to the surface of the giant mitochondria during elongation ^14,18,37^. The testis-specific splice variant of Centrosomin (Cnn), CnnT, was also reported to promote non-centrosomal MT growth on the surface of the mitochondrial derivative of spermatids, and the N-terminal region of Cnn-T was shown to bind efficiently to γ-TuRCs, but there was no data about the canonical γ-TuRC localization after the meiotic stages of spermatogenesis^18,49^. Mutants for the small complex components Grip91 and Grip84 are lethal and have defects in spindle assembly ^21,22^, which indicates their essentiality for the whole fly development. It has been demonstrated that the small complex is the minimal unit needed for MT nucleation ^7^. However, RNAi depletion of the ubiquitously expressed Grip163 results in loss of Grip84 recruitment to the ncMTOC in tracheal cells ^9^.

**Figure 9:**
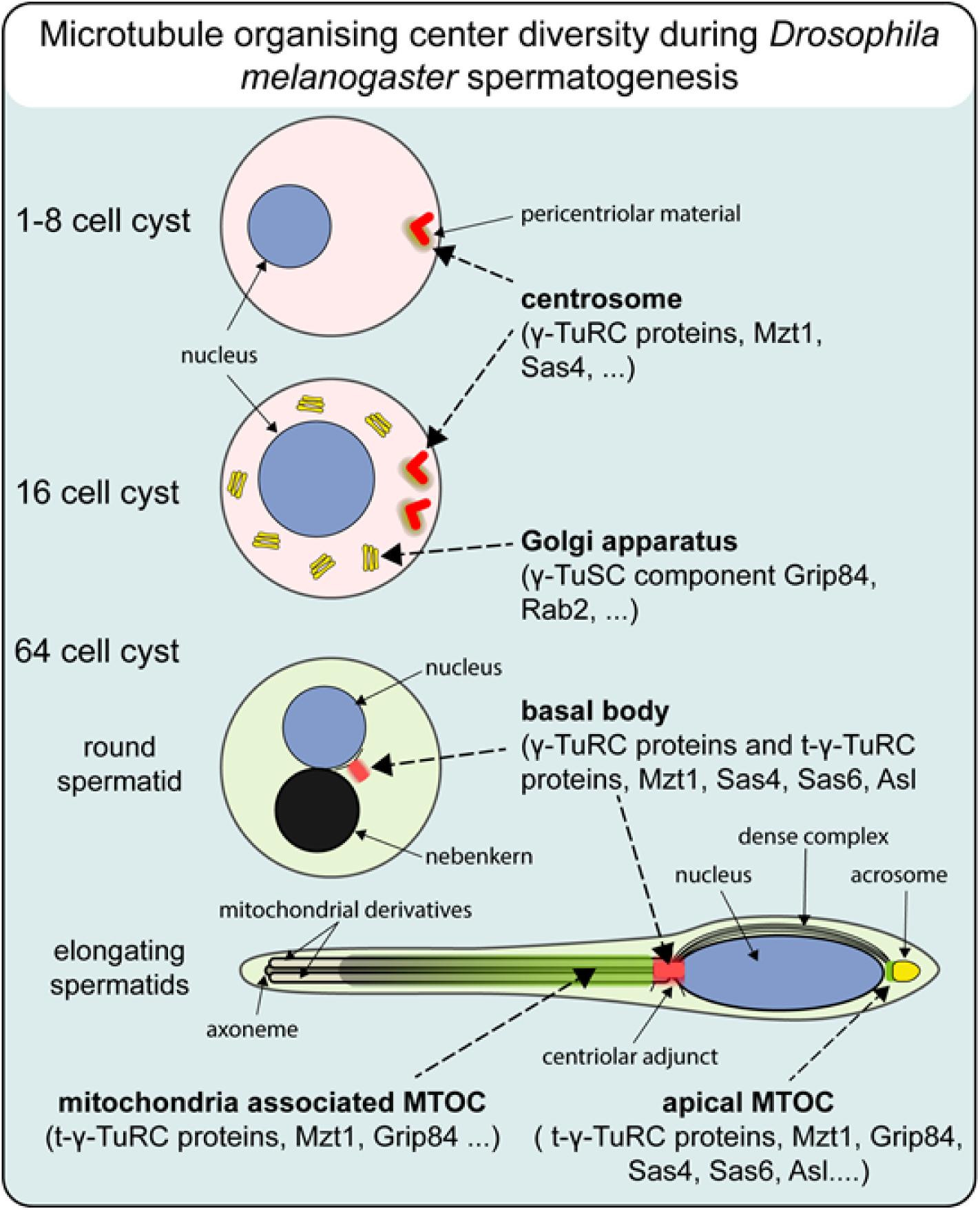
Summary of localization of γ-TuRC/t-γ-TuRC containing MTOCs during different stages of spermatogenesis

Our experiments showed that, in contrast to the essentiality of the γ-TuSC, the γ-TuRC member Grip163 is only essential for fertility, similar to the already characterized γ-TuRC members Grip75 and Grip128 ^23^. From the elongation and basal body stability defects of *Grip163* mutant spermatids, we inferred that the presence of Grip163 protein is essential after meiosis. This was supported by the localization of GFP-Grip163, as we detected its presence in the localization foci of t-γ-TuRC members after meiosis. These results raised the possibility that t-γ-TuRC and the canonical γ-TuRC may be present simultaneously, even as a mixed complex, in the aforementioned possible ncMTOCs (**Figure 9**). This hypothesis was further supported by the fact that Grip84, a member of γ-TuSC, also exhibits an overlapping localization pattern in the late stages of spermatogenesis and the physical interaction between t-Grip84 and the ubiquitously expressed Grip91. Our results strongly suggest that both testis-specific and canonical γ-TuRC members are involved in the formation of post-meiotic ncMTOCs. Although we observed very similar enrichment patterns in testis samples, they were not identical to those of the embryo samples. The variation can be elucidated by the distinct MTOCs found at various developmental stages and in diverse localization sites within the testis, each characterized by unique or shared protein compositions of gamma-tubulin complexes. Nevertheless, the biochemical isolation of these MTOCs poses a significant challenge for future studies. Considering both the *in vivo* localization patterns and the *in vitro* interaction findings, we suggest a versatile composition of gamma-tubulin complex formation occurring at the basal body, on the mitochondrial surface, and at the apical tip of the nucleus in Drosophila spermatids.

Based on the localization and mutant phenotype of canonical Grip163 and testis-specific t-Grips, it can be assumed that the larger amount and diverse composition of gamma-tubulin complex is required for the cellular reorganization of the elongating spermatids. Gamma-tubulin complexes could be essential to the stable attachment of the basal body/centriole adjunct to the sperm nucleus and for the nucleation and stabilization of tubulins contributing to the formation of the nearly 1,8 mm long axoneme and also to the nucleation of the axoneme independent microtubules around the elongating mitochondrial derivatives^13,14,28^.

*In vivo* tagging of Grip84 enabled us to describe the Golgi localization of the γ-TuSC component Grip84 in primary spermatocytes, in addition to the extensively studied centrosomal localization of the Grip84 ^22^. Until now γ-Tubulin containing Golgi outpost-associated MT nucleation sites was only described to regulate distal dendritic branching in *Drosophila* neurons ^50^. Golgi non-centrosomal MT assembly is mediated by the γ-Tubulin-interacting AKAP450 protein, which is anchored by the cis-Golgi-associated GM130 protein. This provides evidence that the Golgi apparatus may become a major site of MT nucleation by acting as a γ-TuRC docking site in mammalian cells by a mechanism similar to that in the centrosome ^51^. Our results suggest that a similar mechanism could be present in the primary spermatocytes. Further experiments are needed to determine exactly which isoform of the Grip84 protein is associated with the Golgi (**Figure 9**).

All three proteins of γ-TuSC (Grip84, Grip91 and γ-Tubulin) interact with the constitutively active, GTP-bound Rab2 protein ^35^, and Rab2 and Grip84 both localize to both the Golgi and the apical end of the nucleus, suggesting a role for γ-TuSC and Rab2 in the formation of the Golgi and lysosome-associated acrosome. Rab2A and Rab2B proteins were shown to be components of the proacrosomal membrane and essential for acrosome formation and male fertility in mice ^36^.

Drosophila Rab2 is an essential gene and recruitment of Rab2 takes place on the membranes of late endosomes, where it controls the fusion of LAMP-containing biosynthetic carriers and lysosomes to late endosomes in somatic cells ^52^. t-ER-Golgi units contribute to the formation of the acroblast, which forms the acrosome by fusing lysosomal structures during nuclear elongation in Drosophila ^53^. The molecular determinants of acrosome formation are hardly known in Drosophila, however, several retrograde tethering complexes were described in the formation of a functional Golgi ribbon in the acroblast (Fári et al 2016). Acrosomal proteins such as Sneaky, Lamp1 and Rab2, as well as the ubiquitous and testis-specific γ-TuRC, its partner proteins (Mzt1, Sas4) and other centrosomal proteins (Asl, Sas6), are all localized to the apical tip of the nucleus. Further biochemical studies are needed to clarify the precise interactions between the aforementioned proteins. Nevertheless, these results raise the possibility of a link between the ncMTOC and acrosome organization, which is present at the dense complex forming perinuclear MTs at the apical end of the nuclei (**Figure 9**).

## Materials and Methods

### Fly stocks, mutants, and fertility test

Flies were maintained on cornmeal agar medium at 25°C in standard laboratory conditions. Fly stocks used in this study were obtained from Bloomington Drosophila Stock Center, except stated otherwise: *w*^1118^ used as wild-type control (3605, BDSC), w[1118];Df(3L)BSC574 (25125 BDSC), P{UASp-YFP.Rab2.Q65L}02 (9760, BDSC), P{DJ-GFP.S}AS1/CyO (5417, BDSC). *bam*-Gal4 testis specific driver ^54^ was used (kindly provided by Helen White-Cooper) to express Rab2 in the germline. The β2-tubulin-GFP line was provided by David M. Glover. GFP-PACT line was obtained from Jordan Raff’s ^29^. Mzt1-GFP, HA-t-Grip91 and t-Grip84-mCh were described before ^14^.

To establish the mutant allele *Grip163^Δ^*^11^ the pCFD5-Grip163-gRNA line was crossed with vas-Cas9 source line to generate deletion in *Grip163*. The deletions were identified by *Grip163* specific PCR and further confirmed by direct sequencing. *Grip163^Δ^*^11^ line was established and used for further analysis. To establish the genomic tagged lines of *Grip84* and *Grip163* pCFD5-Grip84-gRNA and pCFD5-Grip163-gRNA lines were crossed with the vas-Cas9 source lines, w[1118]; PBac{y[+mDint2] GFP[E.3xP3]=vas-Cas9}VK00027 (BL51324), y[1] M{RFP[3xP3.PB] GFP[E.3xP3]=vas-Cas9}ZH-2A w[1118]/FM7c (BL51323) respectively, progeny of this cross were injected with the homology arm carrying pJET1.2 plasmid (Thermo Fisher Scientific). Transgenic candidates were identified by gene specific PCR and confirmed by sequencing of the insertion sites. For fertility tests, individual males were crossed with three w^1118^ virgin females. After 3 days the parents were removed and 14 days after crossing, the hatched progeny was counted in every tube. Experiments were repeated 4 times with > 10 males per condition.

### DNA constructs

To make the genomic GFP-Grip84, and GFP-Grip163 line, annealed oligos of the sgRNAs (ATCCTTGAGCGGCTACCGCT), (GTGCTTCTCCGGTCCAGGCT) for Grip84 and Grip163 respectively were ligated into pCFD5 vector (Addgene, 112645) at Bbs1 restriction site. To establish the homology arms 1534bp upstream, 1529bp downstream, 1515bp upstream and 1738bp downstream of the intended GFP insertion site for Grip84 and Grip163 respectively in addition to GFP were amplified using High fidelity DNA polymerase. The purified PCR products were cloned into pJET1.2 plasmid (Thermo Fisher Scientific) using HiFi DNA Assembly Master Mix (E2621S, NEB) ^55^. GFP sequence was introduced 187bp upstream for Grip84, and 229bp upstream of the Grip163 sgRNA. Stable transgenic lines were established as described in Port et al. ^56^; ^57^.

For Y2H analysis t-Grip84-NT (1-399 aa)^58^ t-Grip84-CT (400-728 aa)^58^, t-Grip128-NT (1-550 aa)^58^, t-Grip128-CT (551-975 aa)^58^ and t-Grip91-NT (1-966 aa)^58^, t-Grip91-CT (967-1932 aa)^58^ were cloned into pGAD424. Full-length of Sas4 (1-901 aa), Grip84 (1-833 aa (Grip84-PD)), Grip75 (1-650 aa) and Grip91-NT (1-545 aa), Grip91-CT (546-917 aa), Grip163-NT (1-685 aa) Grip163-CT (686-1351 aa), Grip128-NT (1-585 aa) Grip128-CT (586-1092 aa) were cloned into pGBT9, using HiFi DNA Assembly Master Mix (E2621S, NEB). To generate GST-tagged bait proteins for the *in vitro* pull-down assay, full-length Grip84, N- and C-terminal parts of t-Grip84, t-Grip91 and t-Grip128 were inserted into the pDEST15 (11802014, Thermo Fisher Scientific) vector by Gateway cloning (11791-020, Thermo Fisher Scientific). Sas4 and Grip91 cDNAs were obtained from the *Drosophila* Gold collection (DGRC) and inserted into the pDEST17 plasmid (11803012, Thermo Fisher Scientific) by Gateway cloning. Oligonucleotide primers used in cloning are listed in **(S. Table 2)**.

### Staining and microscopy

Preparation and staining of testes with formaldehyde or methanol:acetone fixation were performed as described by White-Cooper *et. al* ^59^. Briefly, testes were dissected and fixed in 4% formaldehyde for 25 minutes or semi-squashed using a coverslip on poly-L-lysine coated slides, then frozen with liquid nitrogen and fixed for 5 minutes in ice-cold methanol followed by 5 minutes in acetone. After fixation samples were blocked and antibodies were added, the antibodies used in the study are listed in **(S.Table 1)**. Stained samples were mounted in SlowFade Gold antifade reagent (S36967, Life Technologies). Images were taken using Olympus BX51 fluorescent microscope or Olympus Fluoview Fv10i confocal microscope.

Testes for transmission electron microscopy were dissected and fixed overnight at 4°C in 2% paraformaldehyde and 2% glutaraldehyde. After rinsing in phosphate-buffered saline (PBS), the testes were further fixed for 1 h in 2% OsO_4_, followed by dehydration in increasing ethanol concentrations and rinsing in uranyl acetate and acetone. The samples were embedded in Embed812 (Electron Microscopy Sciences, Hatfield, PA, USA) then ultrathin (70 nm) sections were cut on an Ultracut S ultra-microtome (Leica, Wetzlar, Germany), which were mounted on copper grids. The grids were counterstained with uranyl acetate (Merck, Germany) and lead citrate (Merck, Germany) and were examined and photographed with a JEOL JEM 1400 transmission electron microscope (JEOL, Tokyo, Japan). The images were processed with the TEM Center software (JEOL, Tokyo, Japan).

### Yeast two-hybrid assay

Y2H assays were carried out using the Matchmaker two-hybrid system (K1605, Clontech). One of the baits pGBT9 was cotransformed with one of the preys pGAD424 into PJ69 yeast cells ^60^. Individual colonies were streaked out on yeast synthetic double drop-out plates that lacked tryptophan and leucine (Y0750, Merck), after growing, they were inoculated on double and triple dropout plates lacking (tryptophan, leucine, histidine, and adenine (Y2021, Merck) contains 5-10 mM 3-amino-1, 2, 4-aminotrizole (3-AT) (A8056, Sigma).

### Recombinant protein production

Recombinant GST-tagged bait proteins were expressed, purified and immobilized onto glutathione sepharose beads as described previously ^61^.

### Coupled *in vitro* transcription-translation (IVTT) reaction and *in vitro* binding assay

^35^S-methionine-labelled prey proteins (Grip91, Sas4) were produced *in vitro* using the TnT T7 Quick Coupled IVTT kit (L1170, Promega). The detailed protocol of the GST-IVTT binding assay is described previously ^62^.

### Gel-filtration chromatography and western blotting

Separation of gamma-tubulin-containing complexes were carried out by size exclusion chromatography (SEC) based on the protocol published earlier ^44^ with slight modifications. All steps were conducted at 4°C or on ice unless otherwise stated. Briefly, 25 mg of 0-5 h old dechorionized embryos (expressing GFP-Grip84 or GFP-Grip163) or 500 dissected testes (expressing HA-t-Grip91, t-Grip84-mCh, GFP-Grip84, or GFP-Grip163, respectively) were homogenized in 350 µl HB buffer (50 mM K-HEPES pH 7.6, 75 mM KCl, 1 mM MgCl_2_, 1 mM EGTA, 1 mM DTT, 1 mM PMSF, and EDTA-free protease inhibitor cocktail (PIC, Roche, #11873580001)) using a Dounce homogenizer. Ten strokes with a loose pestle followed by ten strokes with a tight pestle were applied, followed by further homogenization by a plastic pestle in microfuge tubes for 5 min. The samples were centrifuged at 15,000 *xg* for 30 minutes at 4°C. The supernatants were collected and centrifuged again at 21,000 *xg* for 10 minutes at 4°C. Three hundred microliters of clarified lysates were subjected to SEC using a Superose 6 HR 10/30 FPLC column (Cytiva) pre-equilibrated with HB buffer (without DTT, PMSF, and PIC). Forty-one fractions (0.3 ml each) were collected after 7 ml. Ice-cold acetone (2.1 ml) was added to each fraction, the samples were vortexed, and precipitation was carried out at -20°C for 24 hours followed by centrifugation at 17,000 *xg* for 30 minutes at 4°C. The precipitated proteins were resolved in Laemmli sample buffer, boiled for 5 min, and loaded onto SDS-PAGE, followed by western blot analysis using the indicated antibodies: anti-GFP mouse mAb (1:1000, Roche, #11814460001), anti-mCherry rabbit mAb (1:1000, Abcam, #ab213511), anti-gamma-tubulin mouse mAb (1:5000, Sigma, clone GTU-88, #T6557), anti-HA mouse mAb (1:40, Roche, clone 12CA5, #11583816001), and anti-p54/Rpn10 mouse mAb (1:400, ^45^). The following secondary antibodies were used in western blot: goat anti-mouse IgG conjugated to horseradish peroxidase (1:10,000, Dako, #P044701-2) and goat anti-rabbit IgG conjugated to horseradish peroxidase (1:10,000, Dako, #P044801-2).

### Quantification and statistical analysis

Nuclei and nebenkern were counted based on images taken by Olympus BX51. GFP-PACT signals and nebenkern were counted on living samples. GFP-PACT and γ-Tub intensity was measured by FV10-ASW Viewer software (Ver.4.2) on images taken with the same settings, fluorescence intensity was measured in the axis of PACT-GFP signal. Fluorescence intensity were adjusted based on background.

Data analysis and graph production was done using Origin 9.0 (OriginLab Corporation, Northampton, MA, USA). One-way ANOVA followed by Tukey’s HSD test was used to comparisons between WT and mutants. Additional visualisation were made using Python 3.0 seaborn library.

## Acknowledgements

We are grateful to all members of the Sinka lab. We are indebted to David M. Glover, Helen White-Cooper, Sean Munro and Jordan Raff for fly stocks and reagents; for the Drosophila Genomics Resource Center (supported by NIH grant 2P40OD010949) for cDNA clones and to Ferenc Jankovics for critical reading of the manuscript.

---

**Supplementary Table 1.**
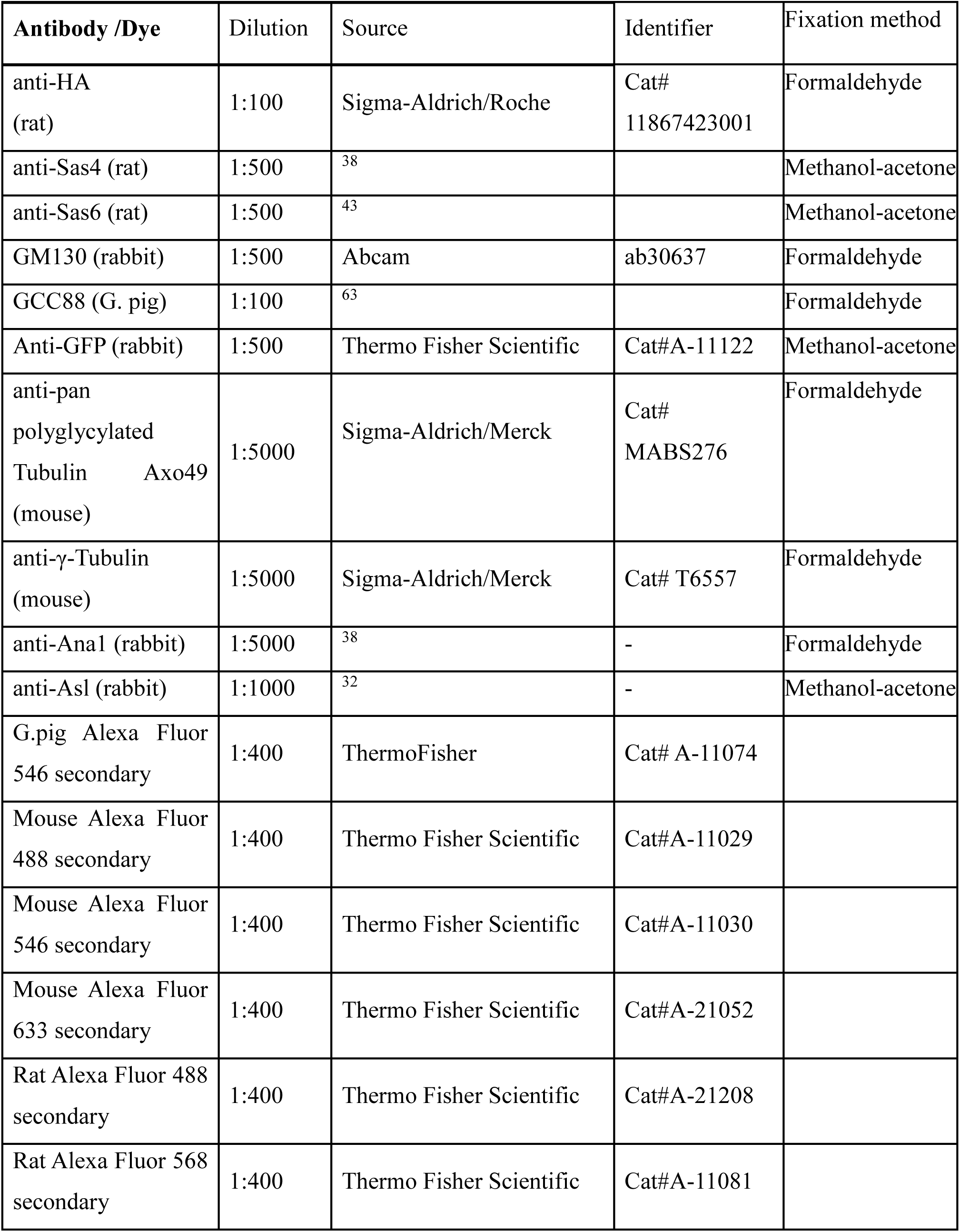
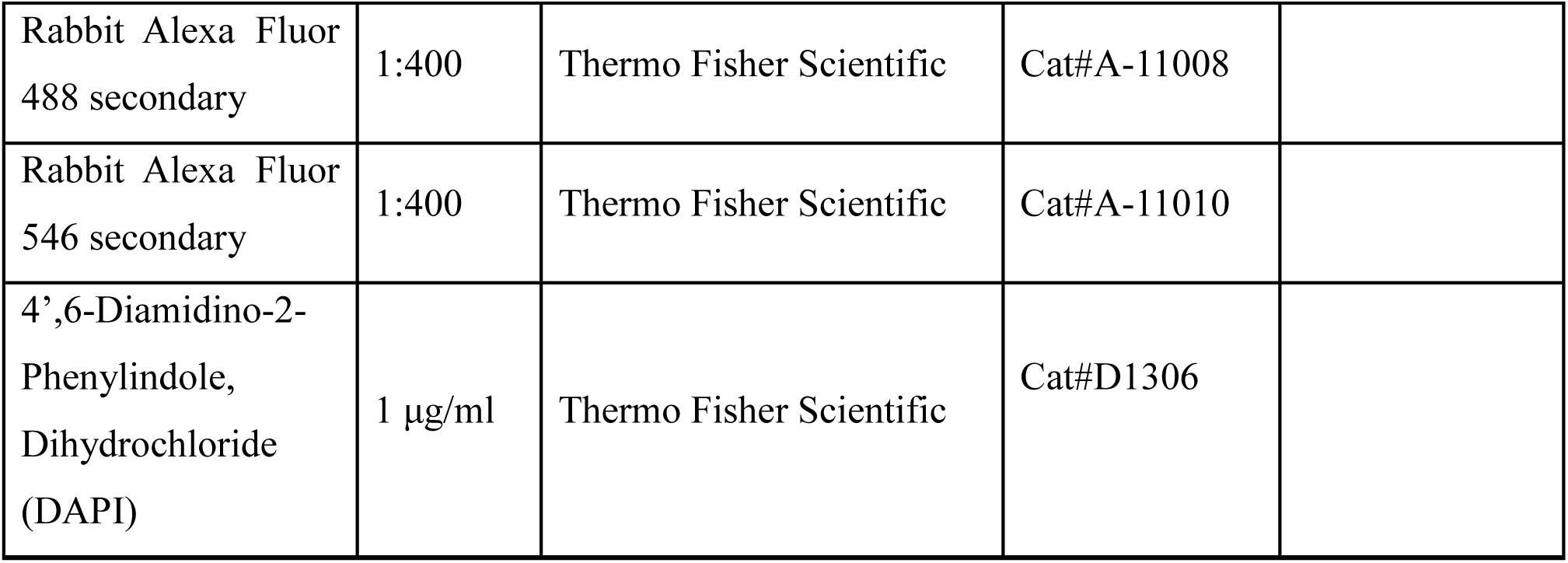
List of antibodies used in immunostaining.

**Supplementary Table 2.**
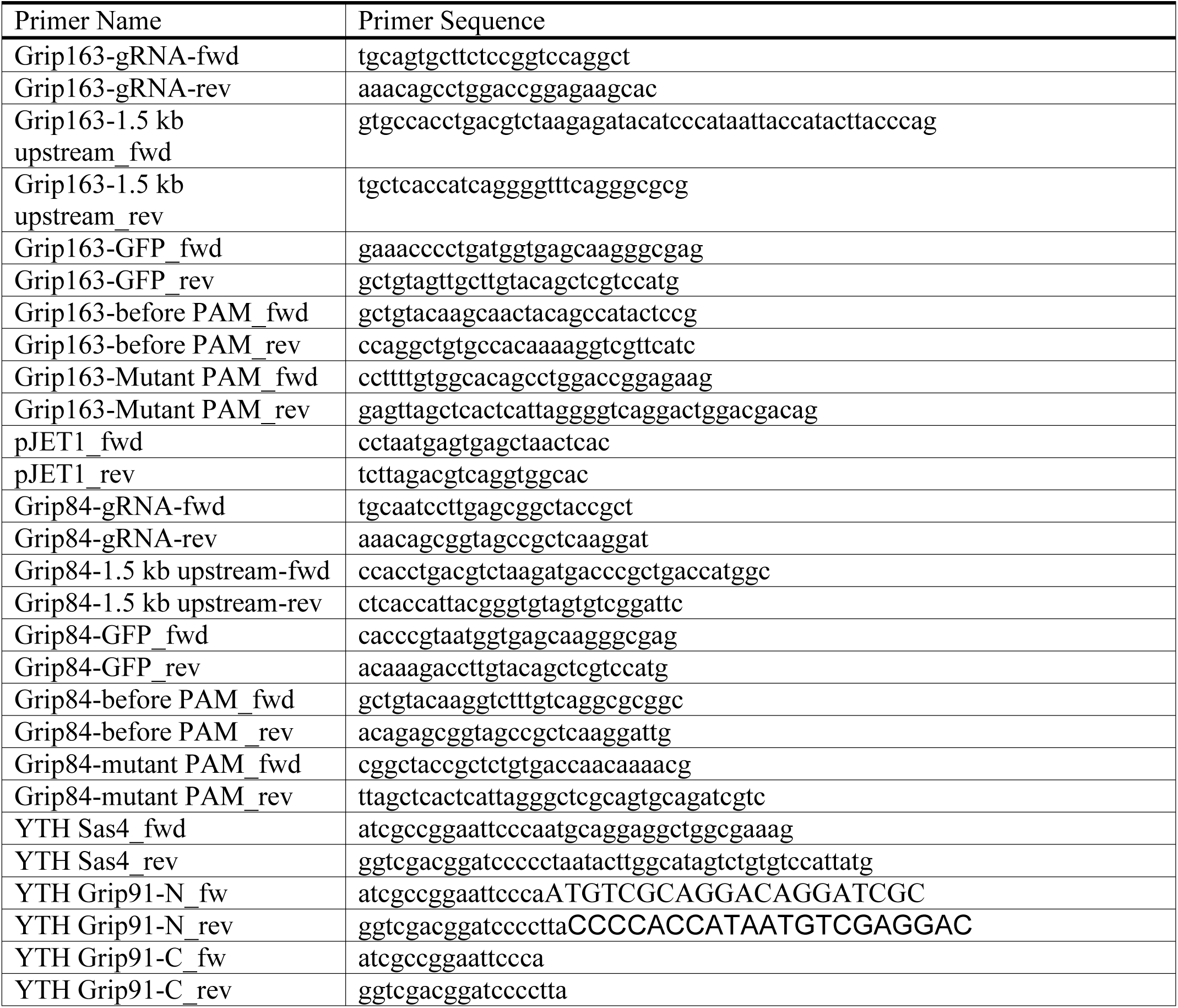
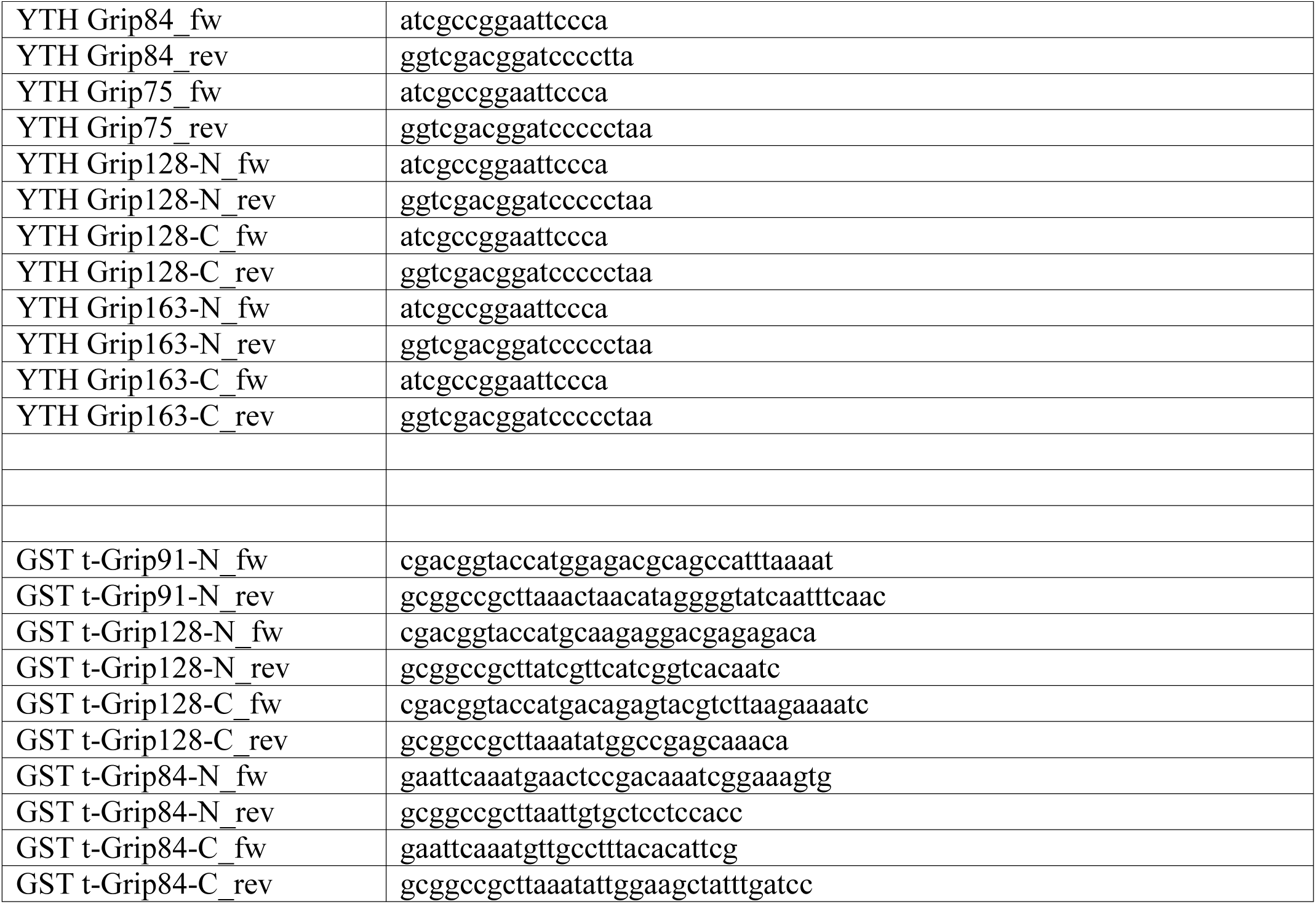
List of oligonucleotide primers uses in this study

**Supplementary Figure 1:**
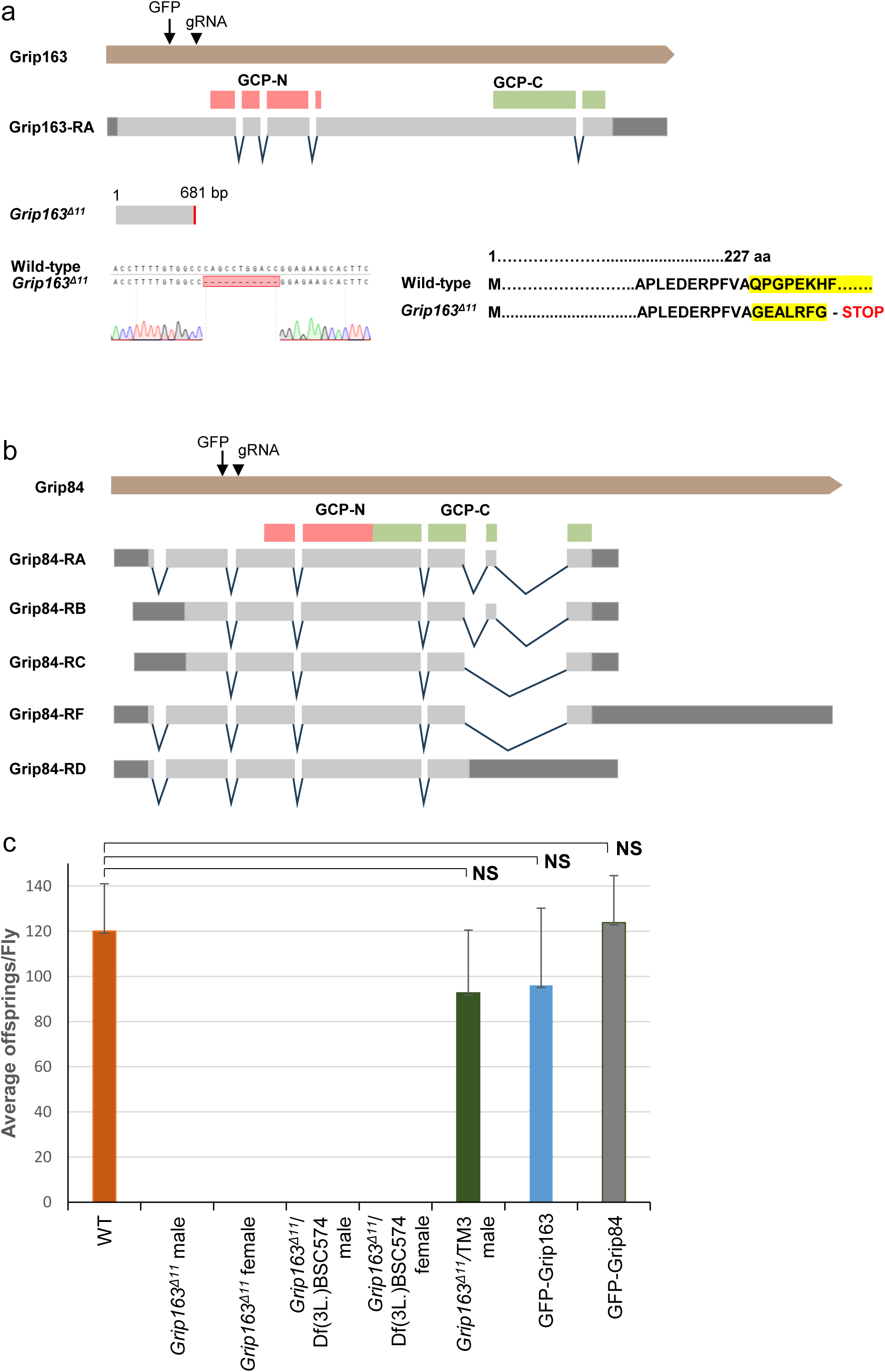
Structural arrangement of *Grip163* and *Grip84* genes and transcripts. **(a)** Gene structure, transcript composition and protein domains (GCP-N, GCP-C) of Grip163. gRNA position (11,997,349..11,997,371 [+]) and the position of the *in vivo* inserted GFP tag (11,997,601 [-]) is labelled. *Grip163^Δ^*^11^ mutant has an 11bp deletion (11,997,369..11,997,359). The DNA and the protein sequences surrounding the deletion are shown. **(b)** Gene structure of Grip84 with the sites of gRNA (19,564,635..19,564,657 [-]) and the GFP tag (19,564,845 [-]). The inserted GFP tag labels all five Grip84 isoforms **(c)** Graph represents the fertility test of WT (n=14), *Grip163^Δ^*^11^ (n=6), *Grip163^Δ^*^11^*/Cyo* (n= 12), GFP-Grip84 (n=12) and GFP-Grip163 (n=15) males. Statistical significance was tested by one-way ANOVA.

**Supplementary Figure 2:**
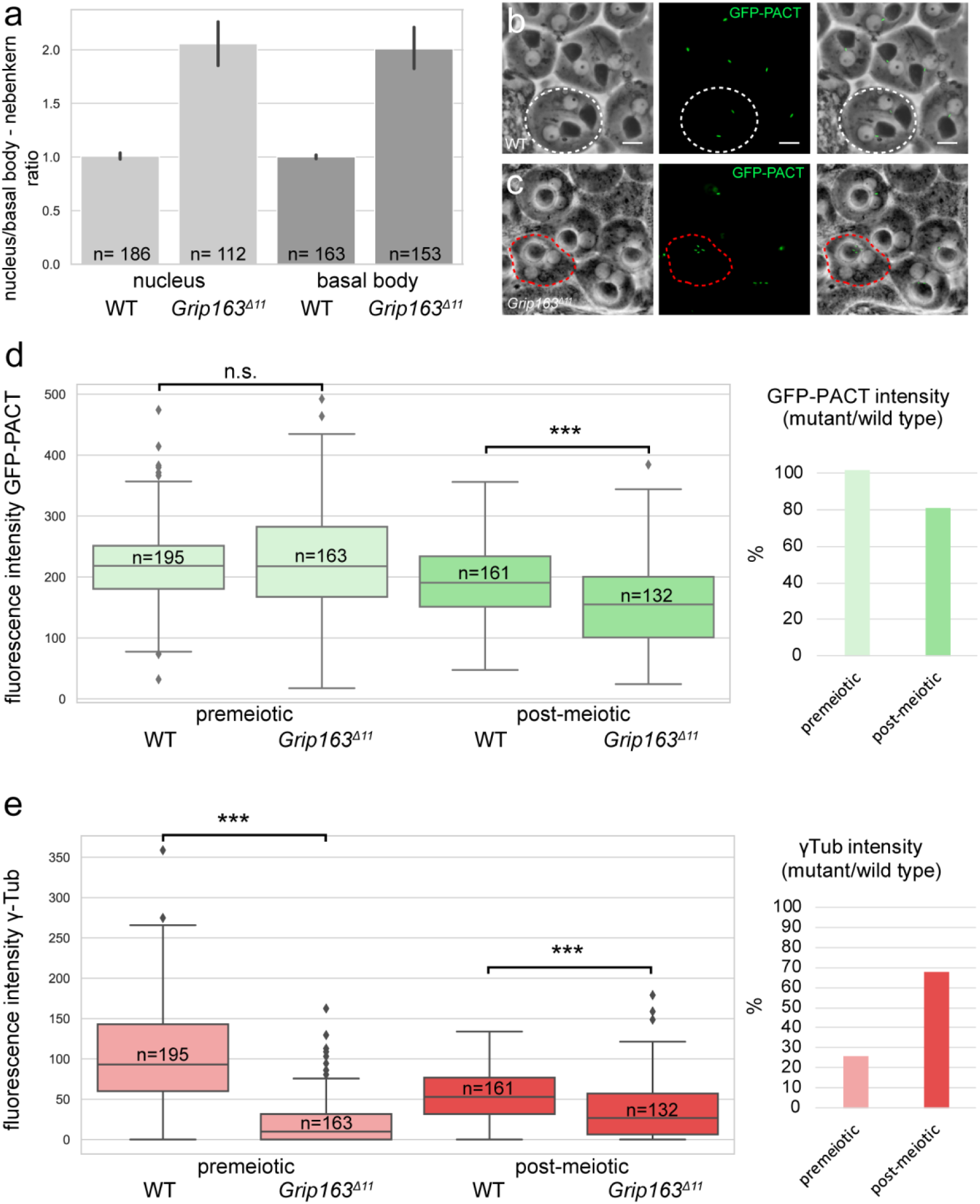
Measurment of distribution of GFP-PACT and γ-Tubulin. (a) Barplot represents the mean values of nucleus/nebenkern and basal body/nebenkern ratios in round spermatids. Error bars show 95% confidence intervals. The number of spermatid cyst fragments is indicated at the base of the columns (each category is represented ∼300 round spermatids). (b, c) Phase contrast images and fluorescent images of GFP-PACT (green) show wild type (b, white dashed line) and Grip163Δ11 mutant round spermatids (c, red dashed line) scale bars represent 10 μm. (d) Boxplot represents fluorescence intensity of GFP-PACT in premeiotic and post-meiotic stages of WT and Grip163Δ11. The number of measurements is indicated on the figure. Barplot indicate the ratio of fluorescence intensity in percentage between the Grip163Δ11 mutant and wild type. (e) Boxplot represents fluorescence intensity of γ-Tub in premeiotic, post-meiotic stages of wild type and Grip163Δ11. The number of measurements is indicated in the figure. The barplot indicates the ratio of fluorescence intensity in percentage between the Grip163Δ11 mutant and wild type.

**Supplementary Figure 3:**
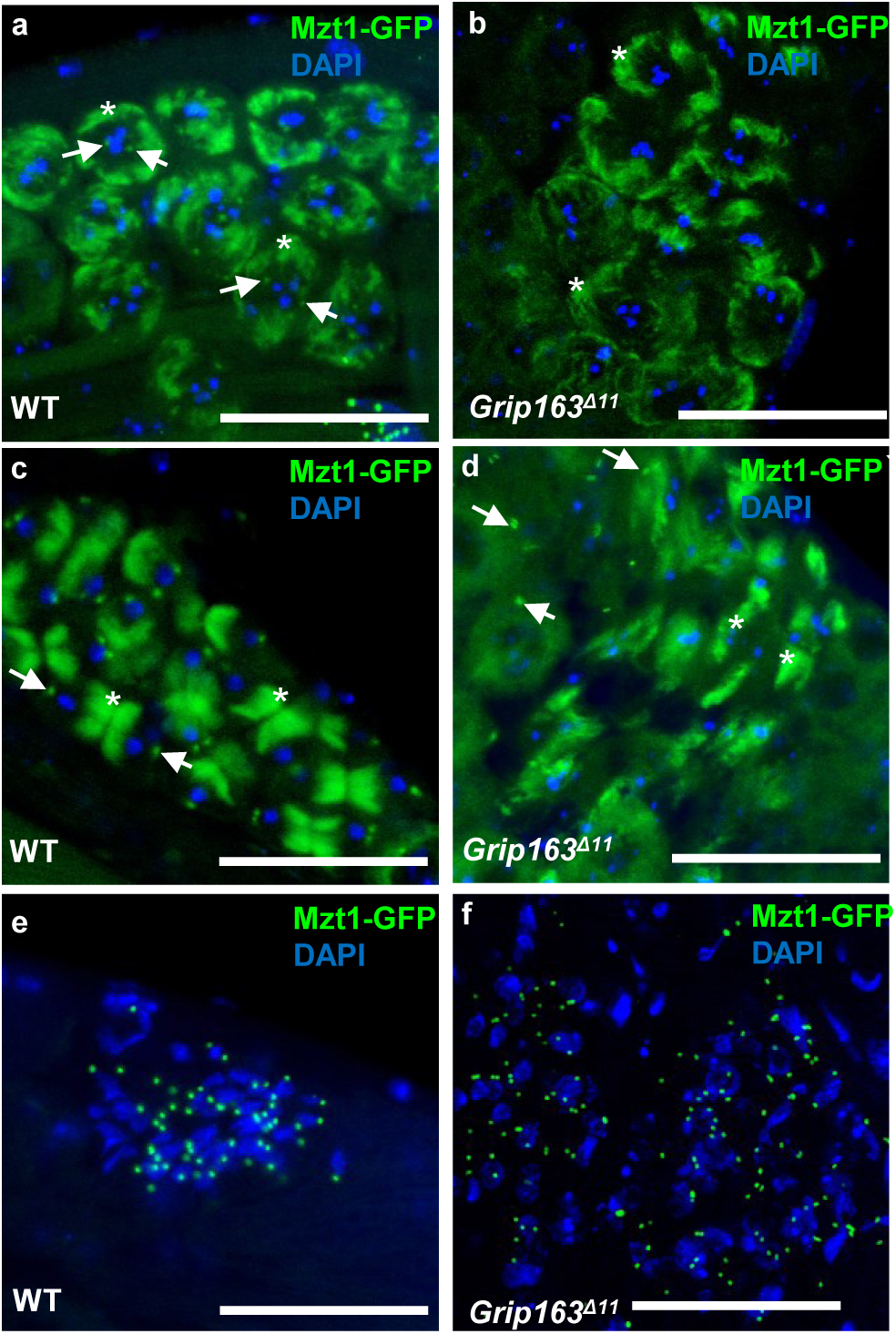
Mzt1-GFP localization in WT and Grip163Δ11 mutant. (a-d) Mzt1-GFP localizes to the basal body (arrow) and mitochondria (asterisk) in WT and to the mitochondria in Grip163Δ11 mutant in dividing meiotic cysts. (e-f) Basal body localization of Mzt1-GFP is present in both in WT and Grip163Δ11 mutant.

**Supplementary Figure 4:**
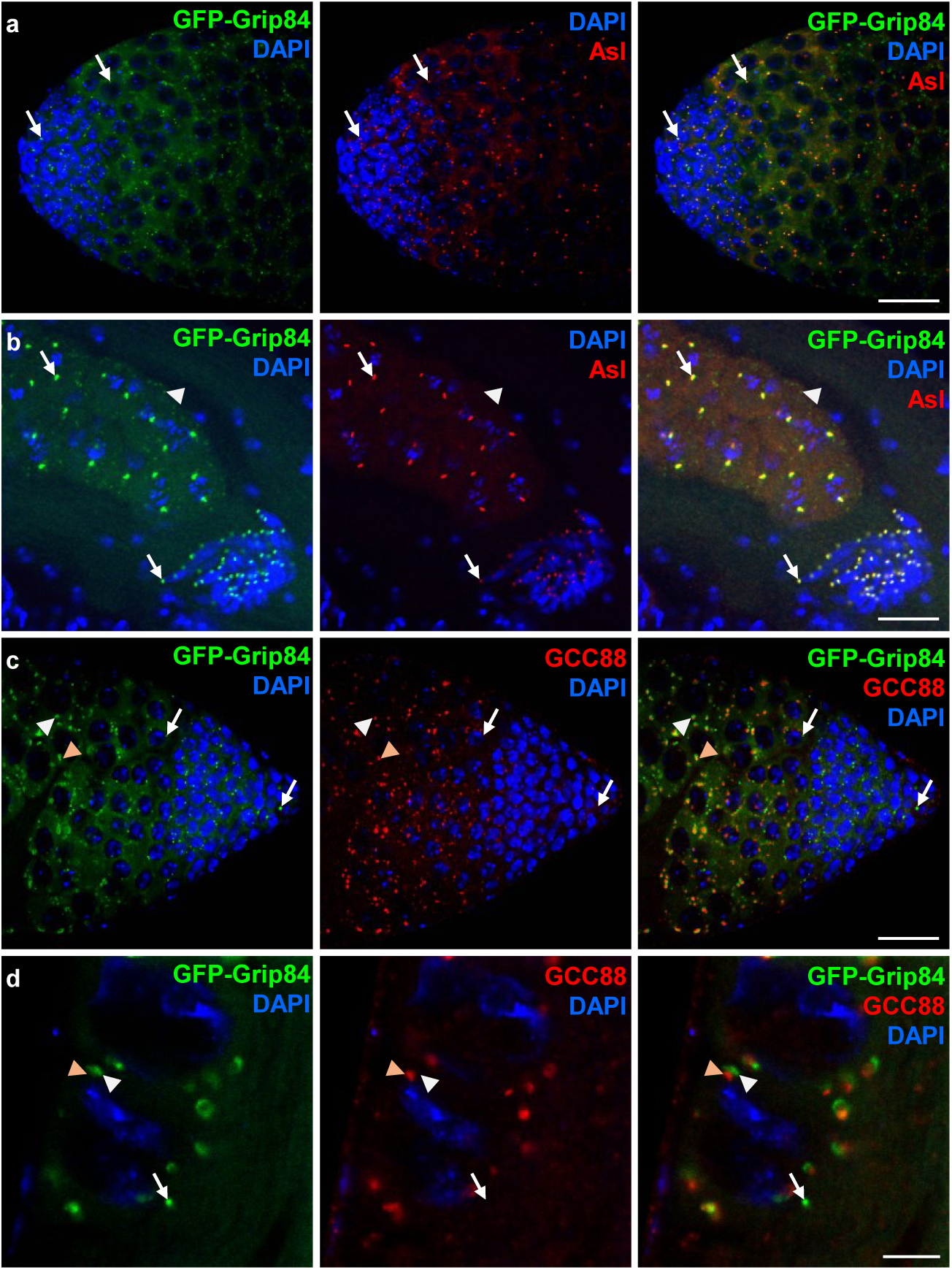
Grip84 localization during spermatogenesis. (a-d) The endogenous tag GFP-Grip84 colocalizes with antibody-stained Asl (red) at the (a) centrosome (arrow) and the (b) basal body (arrow) of the spermatocytes and the spermatids, but not with the Golgi localized GFP-Grip84 (arrowhead). (c, d) GFP-Grip84 localizes adjacent to antibody-stained trans-Golgi marker GCC88 (arrowheads) and to the centrosome (arrow) in spermatocytes. (Scale bars: 20 μm).

**Supplementary Figure 5:**
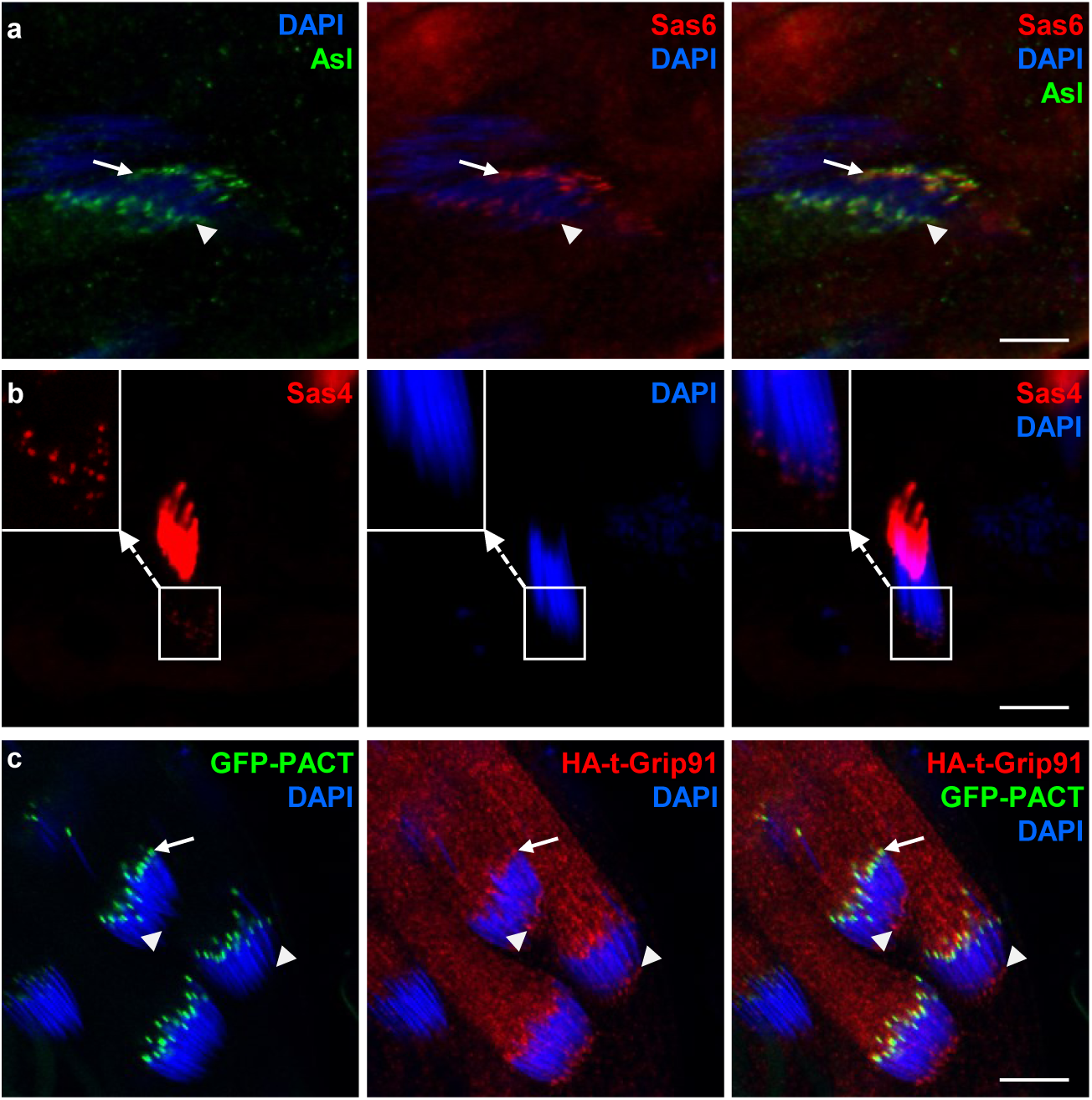
Localization of centrosomal and basal body component proteins in spermatids. **(a, b)** Asl, Sas6 and Sas4 localization to the basal body (arrow) and the apical tip (arrowhead) of elongating spermatids are visualised by immunostaining using gene-specific antibodies. **(c)** GFP-PACT localizes to the basal body (arrows) of the elongating spermatids, while the HA-t-Grip91 localizes to the basal body (arrow) and the apical tip (arrowheads) of the elongating spermatids. (Scale bars: 20 μm)

**Supplementary Figure 6:**
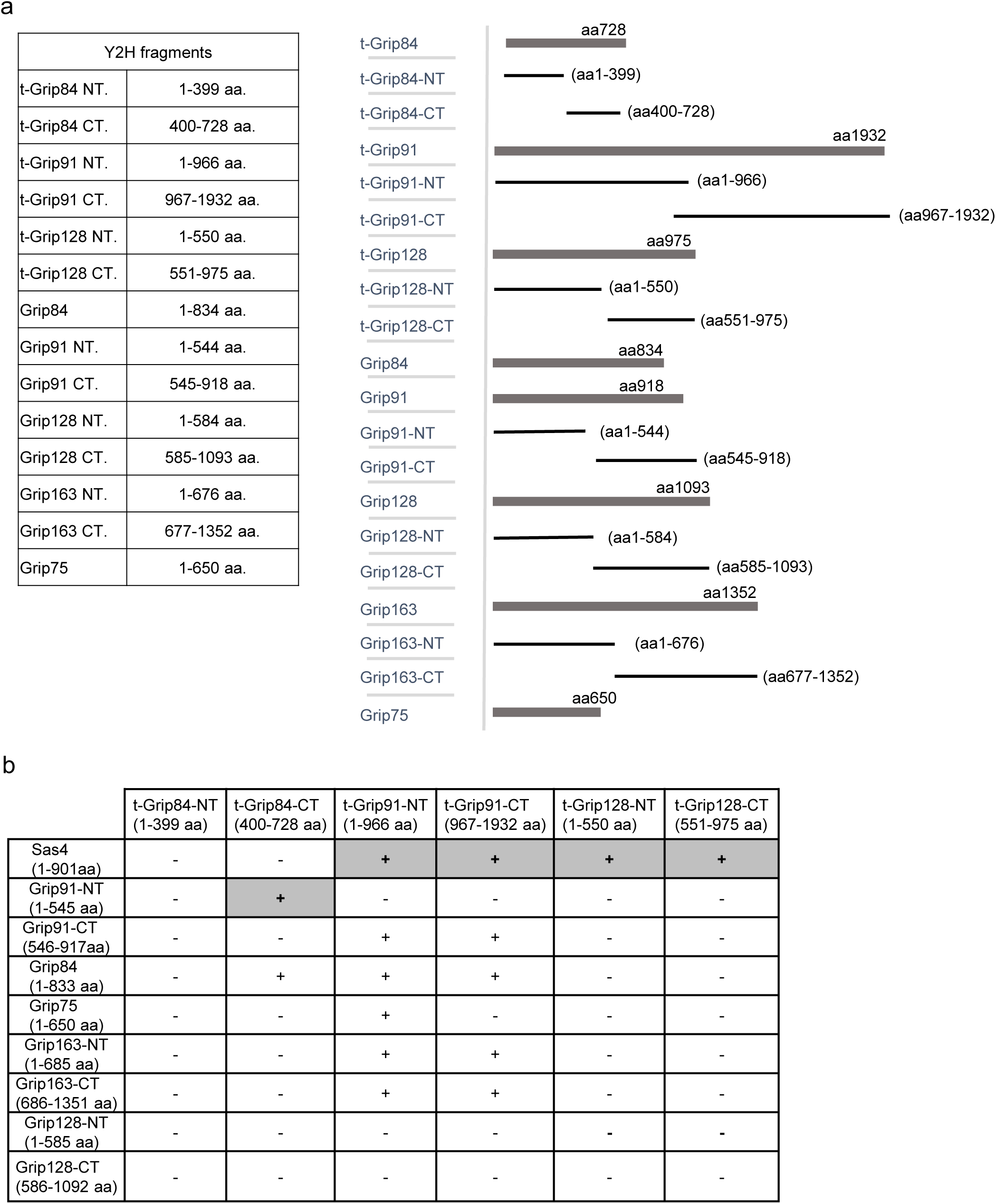
Constructs used in Yeast two-hybrid assay. (a) Truncated fragments of Grip and t-Grip proteins used in Y2H assay are indicated. (b) Summary of the interactions (+) and lack of interactions (-) determined by Y2H analysis. Grey cells show interactions also validated by in vitro protein interaction assay.

**Supplementary Figure 7:**
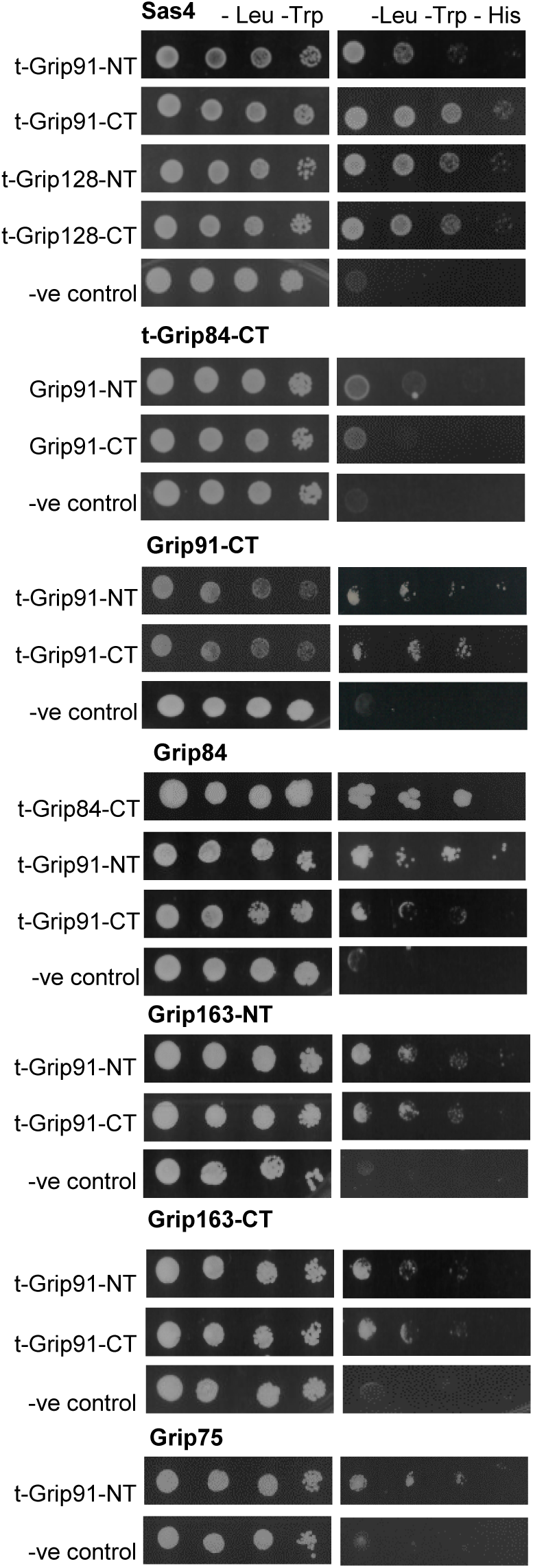
Interactions tested by Yeast Two-Hybrid assay Yeast was plated as 10-fold serial dilutions (left to right) on -Leu, -Trp medium that is selective for the bait and prey plasmids and -Leu, -Trp, -His medium that is selective for the bait, prey, and the interaction between the tested proteins. Negative control (-control) is the empty prey with the corresponding bait vector.

**Supplementary Figure 8:**
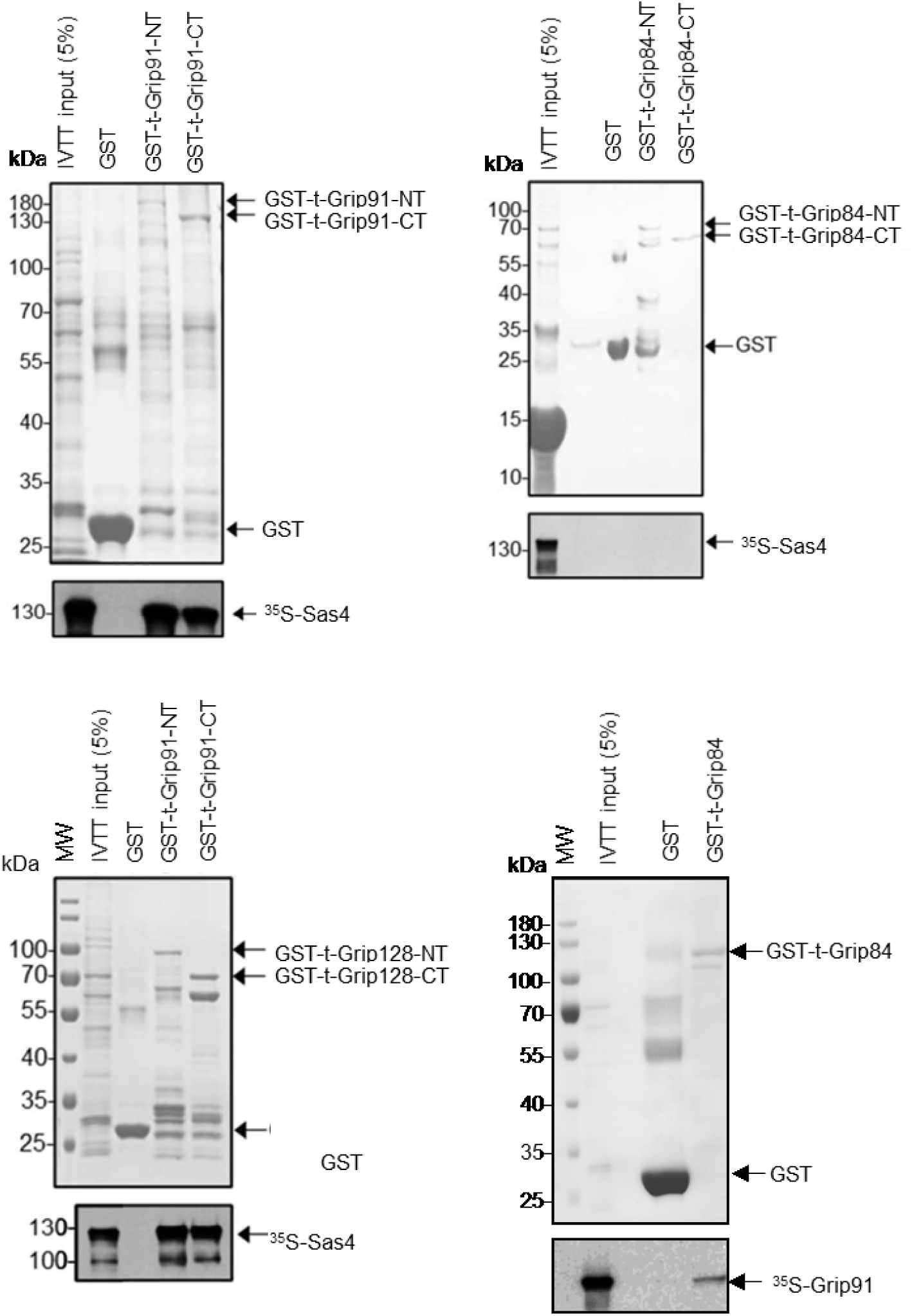
Coomassie-Brilliant Blue stained SDS-PAGE gels (upper panel) show the presence of the purified and immobilized GST-t-Grip91-NT, t-Grip91-CT, GST-t-Grip84-NT, GST-t-Grip84-CT, GST-t-Grip128-NT, GST-t-Grip128-CT or full length of t-Grip84 bait proteins used in the in vitro binding assay. Autoradiographs (lower panel) show the 35S labelled IVTT-produced prey proteins (Sas4 and Grip91).

**Supplementary Figure 9:**
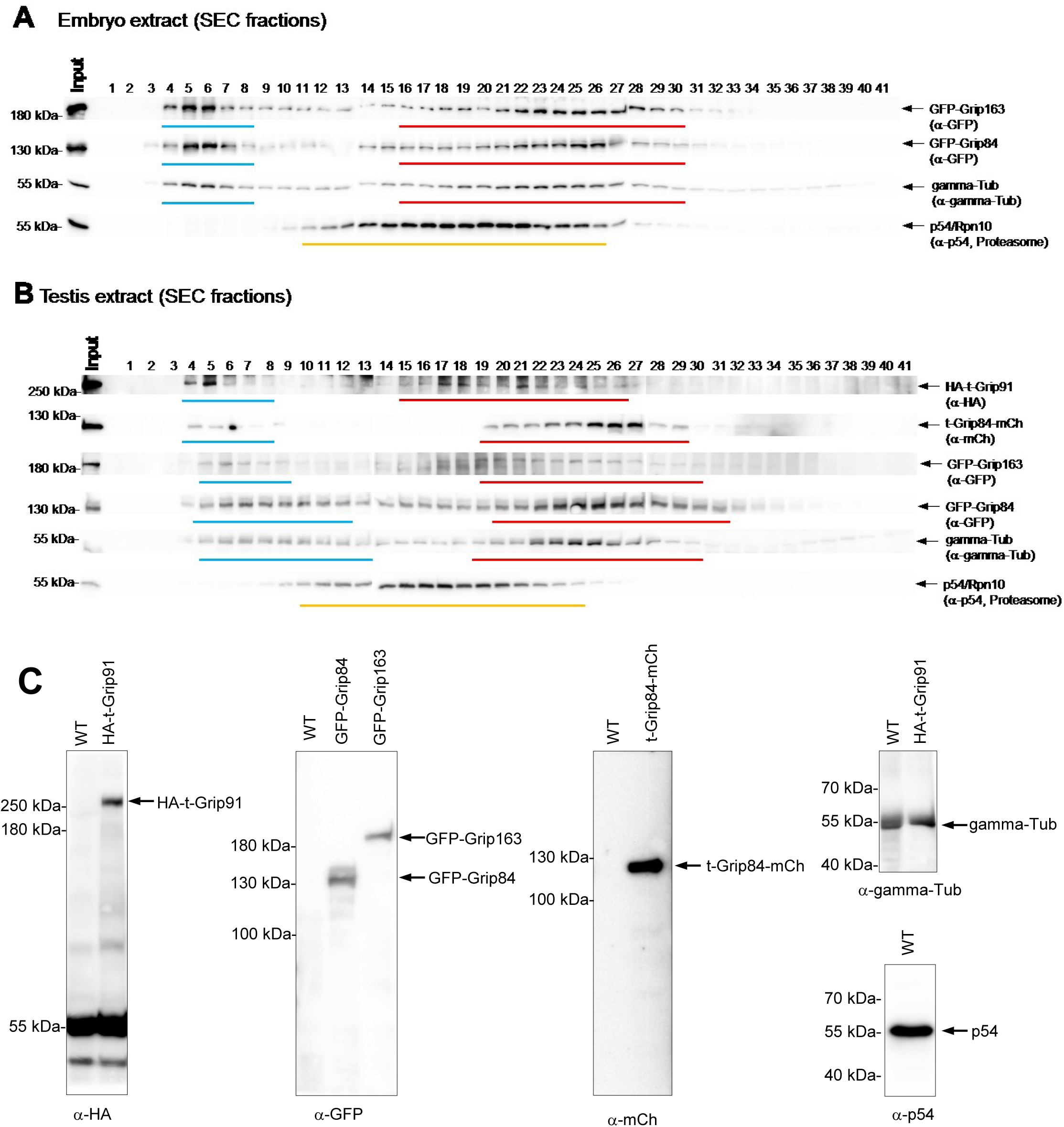
Distinct gamma-tubulin-containing complexes are present in Drosophila testis extracts. The separation of different gamma-tubulin-containing complexes in the embryo (a), as well as testes (b) extracts, was carried out by Superose 6 size-exclusion chromatography (SEC). The fractionation pattern of gamma-tubulin, GFP-Grip163, GFP-Grip84, HA-t-Grip91 and t-Grip84-mCh, which were analysed by western blotting using the indicated antibodies, indicates that differently composed gamma-tubulin complexes may present in testes, compared to embryos. Input refers to the sample loaded onto the column. Blue and red bars represent different complexes with different retention profiles. p54/Rpn10 served as a marker of the 26S proteasome (∼2.5 MDa). (c) Total protein extracts prepared from testes of wild-type (WT) or transgenic flies were analysed by western blotting using the indicated antibodies.

## References

1. Tillery, M., Blake-Hedges, C., Zheng, Y., Buchwalter, R. & Megraw, T. Centrosomal and Non-Centrosomal Microtubule-Organizing Centers (MTOCs) in Drosophila melanogaster. Cells 7, 121 (2018).

2. Desai, A. & Mitchison, T. J. Microtubule polymerization dynamics. Annu. Rev. Cell Dev. Biol. 13, 83–117 (1997).

3. Moritz, M., Braunfeld, M. B., Guénebaut, V., Heuser, J. & Agard, D. A. Structure of the γ-tubulin ring complex: A template for microtubule nucleation. Nat. Cell Biol. 2, 365–370 (2000).

4. Zheng, Y., Wong, M. L., Alberts, B. & Mitchison, T. Nucleation of microtubule assembly by a gamma-tubulin-containing ring complex. Nature 378, 578–583 (1995).

5. Fry, A. M., Sampson, J., Shak, C. & Shackleton, S. Recent advances in pericentriolar material organization: ordered layers and scaffolding gels. F1000Research 6, 1622 (2017).

6. Kollman, J. M., Merdes, A., Mourey, L. & Agard, D. A. Microtubule nucleation by γ-tubulin complexes. Nat. Rev. Mol. Cell Biol. 12, 709–721 (2011).

7. Oegema, K. et al. Characterization of Two Related Drosophila γ-tubulin Complexes that Differ in Their Ability to Nucleate Microtubules. J. Cell Biol. 144, 721–733 (1999).

8. Gunawardane, R. N. et al. Characterization and reconstitution of Drosophila gamma-tubulin ring complex subunits. J. Cell Biol. 151, 1513–1524 (2000).

9. Vérollet, C. et al. Drosophila melanogaster gamma-TuRC is dispensable for targeting gamma-tubulin to the centrosome and microtubule nucleation. J. Cell Biol. 172, 517–528 (2006).

10. Tovey, C. A. & Conduit, P. T. Microtubule nucleation by γ-tubulin complexes and beyond. Essays Biochem. 62, 765–780 (2018).

11. Dhani, D. K. et al. Mzt1/Tam4, a fission yeast MOZART1 homologue, is an essential component of the γ-tubulin complex and directly interacts with GCP3Alp6. Mol. Biol. Cell (2013) doi:10.1091/mbc.e13-05-0253.

12. Masuda, H., Mori, R., Yukawa, M. & Toda, T. Fission yeast MOZART1/Mzt1 is an essential γ-tubulin complex component required for complex recruitment to the microtubule organizing center, but not its assembly. Mol. Biol. Cell (2013) doi:10.1091/mbc.e13-05-0235.

13. Tovey, C. A. et al. γ-TuRC Heterogeneity Revealed by Analysis of Mozart1. Curr. Biol. CB 28, 2314–2323.e6 (2018).

14. Alzyoud, E., Vedelek, V., Réthi-Nagy, Z., Lipinszki, Z. & Sinka, R. Microtubule Organizing Centers Contain Testis-Specific γ-TuRC Proteins in Spermatids of Drosophila. Front. Cell Dev. Biol. 9, 727264 (2021).

15. Tokuyasu, K. T. Dynamics of spermiogenesis in Drosophila melanogaster. VI. Significance of ‘onion’ nebenkern formation. J. Ultrastruct. Res. 53, 93–112 (1975).

16. Avasthi, P. & Marshall, W. F. Stages of ciliogenesis and regulation of ciliary length. Differ. Res. Biol. Divers. 83, S30–42 (2012).

17. Laurinyecz, B. et al. Sperm-Leucylaminopeptidases are required for male fertility as structural components of mitochondrial paracrystalline material in Drosophila melanogaster sperm. PLoS Genet. 15, 1–24 (2019).

18. Chen, J. V., Buchwalter, R. A., Kao, L.-R. & Megraw, T. L. A Splice Variant of Centrosomin Converts Mitochondria to Microtubule-Organizing Centers. Curr. Biol. CB 27, 1928–1940.e6 (2017).

19. Vedelek, V. et al. Analysis of Drosophila melanogaster testis transcriptome 06 Biological Sciences 0604 Genetics. BMC Genomics 19, 1–19 (2018).

20. Wasbrough, E. R. et al. The Drosophila melanogaster sperm proteome-II (DmSP-II). J. Proteomics 73, 2171–2185 (2010).

21. Barbosa, V., Yamamoto, R. R., Henderson, D. S. & Glover, D. M. Mutation of a Drosophila gamma tubulin ring complex subunit encoded by discs degenerate-4 differentially disrupts centrosomal protein localization. Genes Dev. 14, 3126–3139 (2000).

22. Colombié, N. et al. The Drosophila gamma-tubulin small complex subunit Dgrip84 is required for structural and functional integrity of the spindle apparatus. Mol. Biol. Cell 17, 272–282 (2006).

23. Vogt, N., Koch, I., Schwarz, H., Schnorrer, F. & Nüsslein-Volhard, C. The gammaTuRC components Grip75 and Grip128 have an essential microtubule-anchoring function in the Drosophila germline. Dev. Camb. Engl. 133, 3963–3972 (2006).

24. Bärenz, F., Mayilo, D. & Gruss, O. J. Centriolar satellites: Busy orbits around the centrosome. Eur. J. Cell Biol. 90, 983–989 (2011).

25. Gupta, H. et al. SAS-6 Association with γ-Tubulin Ring Complex Is Required for Centriole Duplication in Human Cells. Curr. Biol. 30, 2395–2403.e4 (2020).

26. Fabian, L. & Brill, J. A. Big things come from little packages Drosophila spermiogenesis. 1– 16 (2012).

27. Bre, M. H. et al. Axonemal tubulin polyglycylation probed with two monoclonal antibodies: widespread evolutionary distribution, appearance during spermatozoan maturation and possible function in motility. J. Cell Sci. 109, 727–738 (1996).

28. Noguchi, T., Koizumi, M. & Hayashi, S. Sustained elongation of sperm tail promoted by local remodeling of giant mitochondria in Drosophila. Curr. Biol. 21, 805–814 (2011).

29. Martinez-Campos, M., Basto, R., Baker, J., Kernan, M. & Raff, J. W. The Drosophila pericentrin-like protein is essential for cilia/flagella function, but appears to be dispensable for mitosis. J. Cell Biol. 165, 673–683 (2004).

30. Zupa, E., Liu, P., Würtz, M., Schiebel, E. & Pfeffer, S. The structure of the γ-TuRC: a 25-years-old molecular puzzle. Curr. Opin. Struct. Biol. 66, 15–21 (2021).

31. Bourbon, H.-M. et al. A P-insertion screen identifying novel X-linked essential genes in Drosophila. Mech. Dev. 110, 71–83 (2002).

32. Dzhindzhev, N. S. et al. Asterless is a scaffold for the onset of centriole assembly. Nature 467, 714–718 (2010).

33. Allan, V. J., Thompson, H. M. & McNiven, M. A. Motoring around the Golgi. Nat. Cell Biol. 4, E236–E242 (2002).

34. Kondylis, V. & Rabouille, C. The Golgi apparatus: Lessons from Drosophila. FEBS Lett. 583, 3827–3838 (2009).

35. Gillingham, A. K., Sinka, R., Torres, I. L., Lilley, K. S. & Munro, S. Toward a Comprehensive Map of the Effectors of Rab GTPases. Dev. Cell 31, 358 (2014).

36. Morohoshi, A. et al. FAM71F1 binds to RAB2A and RAB2B and is essential for acrosome formation and male fertility in mice. Development 148, dev199644 (2021).

37. Riparbelli, M. G., Persico, V. & Callaini, G. The Microtubule Cytoskeleton during the Early Drosophila Spermiogenesis. Cells 9, 1–18 (2020).

38. Fu, J. et al. Conserved molecular interactions in centriole-to-centrosome conversion. Nat. Cell Biol. 18, 87–99 (2016).

39. Gopalakrishnan, J. et al. Tubulin nucleotide status controls Sas-4-dependent pericentriolar material recruitment. Nat. Cell Biol. 14, 865–873 (2012).

40. Galletta, B. J. et al. A centrosome interactome provides insight into organelle assembly and reveals a non-duplication role for Plk4. Nat. Commun. 7, 12476 (2016).

41. Gopalakrishnan, J. et al. Sas-4 provides a scaffold for cytoplasmic complexes and tethers them in a centrosome. Nat. Commun. 2, 359 (2011).

42. Guruharsha, K. G. et al. A Protein Complex Network of Drosophila melanogaster. Cell 147, 690–703 (2011).

43. Dzhindzhev, N. S. et al. Plk4 phosphorylates Ana2 to trigger Sas6 recruitment and procentriole formation. Curr. Biol. CB 24, 2526–2532 (2014).

44. Moritz, M., Zheng, Y., Alberts, B. M. & Oegema, K. Recruitment of the gamma-tubulin ring complex to Drosophila salt-stripped centrosome scaffolds. J. Cell Biol. 142, 775–786 (1998).

45. Kurucz, E. et al. Assembly of the Drosophila 26 S proteasome is accompanied by extensive subunit rearrangements. Biochem. J. 365, 527 (2002).

46. Efimov, A. et al. Asymmetric CLASP-dependent nucleation of non-centrosomal microtubules at the trans-Golgi network. Dev. Cell 12, 917–930 (2007).

47. Becker, R. et al. Myogenin controls via AKAP6 non-centrosomal microtubule-organizing center formation at the nuclear envelope. eLife 10, e65672 (2021).

48. Brodu, V., Baffet, A. D., Le Droguen, P.-M., Casanova, J. & Guichet, A. A developmentally regulated two-step process generates a noncentrosomal microtubule network in Drosophila tracheal cells. Dev. Cell 18, 790–801 (2010).

49. Tovey, C. A. et al. Autoinhibition of Cnn binding to γ-TuRCs prevents ectopic microtubule nucleation and cell division defects. J. Cell Biol. 220, e202010020 (2021).

50. Mukherjee, A., Brooks, P. S., Bernard, F., Guichet, A. & Conduit, P. T. Microtubules originate asymmetrically at the somatic golgi and are guided via Kinesin2 to maintain polarity within neurons. eLife 9, e58943 (2020).

51. Rivero, S., Cardenas, J., Bornens, M. & Rios, R. M. Microtubule nucleation at the cis-side of the Golgi apparatus requires AKAP450 and GM130. EMBO J. 28, 1016–1028 (2009).

52. Lund, V. K., Madsen, K. L. & Kjaerulff, O. Drosophila Rab2 controls endosome-lysosome fusion and LAMP delivery to late endosomes. Autophagy 14, 1520–1542 (2018).

53. Fári, K., Takács, S., Ungár, D. & Sinka, R. The role of acroblast formation during *Drosophila* spermatogenesis. Biol. Open 5, 1102–1110 (2016).

54. Chen, D. & McKearin, D. M. A discrete transcriptional silencer in the bam gene determines asymmetric division of the Drosophila germline stem cell. Dev. Camb. Engl. 130, 1159–1170 (2003).

55. Gibson, D. G. et al. Enzymatic assembly of DNA molecules up to several hundred kilobases. Nat. Methods 6, 343–345 (2009).

56. Port, F. & Bullock, S. L. Creating Heritable Mutations in Drosophila with CRISPR-Cas9. In Methods in Molecular Biology vol. 1478 145–160 (2016).

57. Port, F., Chen, H. M., Lee, T. & Bullock, S. L. Optimized CRISPR/Cas tools for efficient germline and somatic genome engineering in Drosophila. Proc. Natl. Acad. Sci. U. S. A. 111, (2014).

58. Alzyoud, E., Vedelek, V., Réthi-Nagy, Z., Lipinszki, Z. & Sinka, R. Microtubule Organizing Centers Contain Testis-Specific γ-TuRC Proteins in Spermatids of Drosophila. Front. Cell Dev. Biol. 9, (2021).

59. White-cooper, H. Analysis of Meiosis and Morphogenesis. Drosoph. Cytogenet. Protoc. 247, 468 (2004).

60. James, P., Halladay, J. & Craig, E. A. Genomic Libraries and a Host Strain Designed for Highly Efficient Two-Hybrid Selection in Yeast. Genetics 144, 1425–1436 (1996).

61. Karman, Z. et al. Novel perspectives of target-binding by the evolutionarily conserved PP4 phosphatase. Open Biol. 10, 200343 (2020).

62. Réthi-Nagy, Z., Ábrahám, E. & Lipinszki, Z. GST-IVTT pull-down: a fast and versatile in vitro method for validating and mapping protein-protein interactions. FEBS Open Bio 12, 1988–1995 (2022).

63. Jones, R. B. et al. Golgi coiled-coil proteins contain multiple binding sites for rab family G proteins. J. Cell Biol. 183, 607–615 (2008).

